# Transgenic female mice producing *trans* 10, *cis* 12-conjugated linoleic acid present excessive prostaglandin E2, adrenaline, corticosterone, glucagon, and FGF21

**DOI:** 10.1101/2023.04.21.536206

**Authors:** Yu Rao, Lu-Wen Liang, Mei-Juan Li, Yang-Yang Wang, Bao-Zhu Wang, Ke-Mian Gou

**Author notes:** These authors contributed equally to this work (Yu Rao, Lu-Wen Liang, Mei-Juan Li). Corresponding author: Kemian Gou, College of Veterinary Medicine, Yangzhou University, Wenhui South Road, Yangzhou 225009, China.

## Abstract

Dietary *trans* 10, *cis* 12-conjugated linoleic acid (t10c12-CLA) is a potential candidate in anti-obesity trials. A transgenic mouse was previously successfully established to determine the anti-obesity properties of t10c12-CLA in male mice that could produce endogenous t10c12-CLA. To test whether there is a different impact of t10c12-CLA on lipid metabolism in both sexes, this study investigated the adiposity and metabolic profiles of female Pai mice that exhibited a dose-dependent expression of foreign Pai gene and a shift of t10c12-CLA content in tested tissues. Compared to their gender-match wild-type littermates, Pai mice had no fat reduction but exhibited enhanced lipolysis and thermogenesis by phosphorylated hormone-sensitive lipase and up-regulating uncoupling proteins in brown adipose tissue. Simultaneously, Pai mice showed hepatic steatosis and hypertriglyceridemia by decreasing gene expression involved in lipid and glucose metabolism. Further investigations revealed that t10c10-CLA induced excessive prostaglandin E2, adrenaline, corticosterone, glucagon and inflammatory factors in a dose-dependent manner, resulting in less heat release and oxygen consumption in Pai mice. Moreover, fibroblast growth factor 21 overproduction only in monoallelic Pai/wt mice indicates that it was sensitive to low doses of t10c12-CLA. These results suggest that chronic t10c12-CLA has system-wide effects on female health via synergistic actions of various hormones.

## 1. INTRODUCTION

*Trans* 10, *cis* 12-conjugated linoleic acid (t10c12-CLA) occurs naturally in small amounts in beef and dairy products. Its popular health benefit is to reduce body fat based on energy balancing by increasing energy expenditure via browning of white adipose tissue (WAT) ^1,2^ or uncoupling protein (UCP) 1 & 2-mediated thermogenesis in brown adipose tissue (BAT) ^3^, promoting biosynthesis of N-lactoyl-phenylalanine via the up-regulation of cytosolic nonspecific dipeptidase 2 in WAT ^4^, or increasing intramuscular fat ^5^. In addition, t10c12-CLA has other potential health influences, including protecting against atherosclerosis by modulating the macrophage phenotype ^6^, promoting beneficial effects on cholesterol efflux ^7^, reducing oxidative stress ^8^, changing intestinal microbiota ^9^ and metabolite profiles ^5^, even down-regulating Alzheimer’s hallmarks in an aluminium mouse through a Nuclear factor erythroid 2–related factor 2-mediated adaptive response and increasing brain glucose transporter levels ^10^.

Although the potential health benefits of dietary t10c12-CLA have been extensively studied in mice, other studies also indicated its side effects manifested as fatty liver in male ^9,11^ or female mice ^12-16^. At the same time, a study in mice treated with dietary t10c12-CLA saw a reduction in fat mass in the first two weeks, which interestingly rebounded during the following 3-5 weeks due to a massive accumulation of lipids in the liver ^11^. That being said, the potential health issues of t10c12-CLA, namely its long-term negative effects on the body, have not been fully clarified.

In our previous study, we established a transgenic mouse that produces t10c12-CLA continuously after inserting the foreign gene encoding the *Propionibacterium acnes* isomerase (PAI) into the Rosa26 locus ^3^ for this allele is a genomic safe-harbour site integrating transgene constructs to achieve ubiquitous gene expression in mice ^17^. This Pai knock-in mouse could be used to elucidate the enduring influence of t10c12-CLA on overall health in a novel way. Among its male offspring, the phenotyping of monoallelic Pai/wt mice suggested that the low doses of t10c12-CLA possibly played an active role in lipid metabolism by stimulating the fibroblast growth factor 21 (FGF21) secretion. In contrast, the phenotyping of biallelic Pai/Pai mice indicated that the high doses of t10c12-CLA induced fat reduction by promoting energy expenditure via UCP1/2-mediated BAT thermogenesis^3^.

To test whether there is a different impact of t10c12-CLA’s on lipid metabolism in both sexes, the current study continued investigating female adiposity characteristics and metabolic profiles using Pai mice. In contrast to their male counterparts, the findings of this study demonstrated that Pai female mice did not experience changes in body fat yet experienced excessive prostaglandin E2 (PGE2), corticosterone, glucagon, and adrenaline in a dose-dependent manner, and resulted in inflammation and less heat release. Moreover, Pai/wt female mice further confirmed that the FGF21 overproduction was sensitive to the low doses of t10c12-CLA. These results suggest that chronic t10c12-CLA may affect lipid metabolism by excessive lipid-related hormones synergistically in female mammals. The probable hormone-mediated metabolic syndrome must be considered carefully in t10c12-CLA practice.

## 2. MATERIALS AND METHODS

### 2.1. Mice and diet

The transgenic founder from a hybrid embryo of DBA/2 and C57BL/6J was backcrossed to C57BL/6J mice for more than eight generations, and then the monoallelic Pai/wt matings were used to produce the Pai/Pai, Pai/wt, and wild-type (wt) offspring for this study. Genotyping was performed using genomic DNA from tail tissues, with the presence of the Pai gene being assayed by amplifying the 514-bp Pai fragments (forward: 5-taaccatgttcatgccttcttc-3; reverse: 5-caccttgttgtagtgtccgttt-3) and the wt Rosa26 sequence being assayed by amplifying the 605-bp fragments that spanned the insertional site of the Rosa26 locus (forward: 5-ccaaagtcgctctgagttgttatcagt-3; reverse: 5-ggagcgggagaaatggatatgaag-3). PCR amplification was performed for 30 cycles at 94°C for 30 sec, 57°C for 30 sec, and 72°C for 40 sec.

Mice of the above three genotypes aged 11-15 weeks were used for analysis in this study unless otherwise specified. All animal experiments were conducted under the guidance of the Committee for Experimental Animals of Yangzhou University. The animal study protocol was approved by the Ethics Committee of Yangzhou University (protocol code NSFC2020-SYXY-20 and dated 25 March 2020). The mice were housed in a light-controlled room (12L:12D, lights on at 0700 h) at 22-23°C and fed ad libitum with a standard diet containing 10% kcal% fat. Fresh diets were prepared every month following formula No. D12450H of OpenSource DIETS^TM^ (Research Diets Inc., NJ, USA) under sterile conditions. The composition of the D12450H and the sources of the food-grade ingredients used in the diets are provided in Supplementary Table S1.

### 2.2. Real-time PCR

All real-time PCRs were carried out in 96-well plates using ChamQ SYBR qPCR Master Mix kit (Nanjing Vazyme Biotech Ltd., China) in the ABI Prism 7500 Sequence Detection System (Applied Biosystems, USA) at 95°C, 2 min, one cycle, followed by 40 cycles of 95°C for 5 sec and 60°C for 32 sec. Each sample was run in triplicate. The relative transcriptional level of the target gene was normalised to one of the endogenous gene expressions of glyceraldehyde-3-phosphate dehydrogenase (Gapdh), ApoB, or 36B4, using the method of 2^-ΔΔCt^. The primer sequences are listed in Supplementary Table S2.

### 2.3. Gas chromatography

The gas chromatography procedure followed a modified method described by Jenkins^18^. A two-step trans-esterification process involving sodium methoxide followed by shorter methanolic HCl was used to methylate each organ/tissue homogenised by grinding in liquid nitrogen. Fatty acid methyl esters were separated on an HP-88 fused-silica capillary column (60 m X 0.25 mm i.d., 0.2-µm film thickness, J & W 112-88A7, Agilent Technologies, USA) and quantified using a fully automated 7890 Network GC System with a flame-ionization detector (Agilent). The program settings followed Jenkins ^18^. C19:0 was used as the internal standard, and the peaks were identified by comparing them with fatty acid standards (Sigma, 47885U and O5632). The area percentage of all resolved peaks was analysed using GC ChemStation Software (Agilent).

### 2.4. Blood parameters measurements

Blood samples were collected from mice fed ad libitum unless otherwise stated. Fresh blood from the tail veins was used to measure circulating glucose levels using a handheld glucose monitor (Accu-Chek^®^ Performa Blood Glucose Meter, Roche). Heparin-treated blood from the tail vein was used to respectively measure the plasma triglycerides (TGs) or high-density lipoprotein (HDL) using a handheld cholesterol monitor (On-Call^®^ CCM-111 Blood Cholesterol System, Aikang Biotech Co. Ltd., China). Blood samples from the submandibular vein of conscious mice were collected during the first 4 to 5 hours of the light phase and used to measure serum corticosterone levels using the Cort ELISA kit (Ruixin Biotech Co. Ltd., China). Serum samples from the orbital sinus vein of anaesthetic mice treated with 1.25% tribromoethanol (T48402, Simga; 150 mg/kg body weight) were used to measure circulating total cholesterol (TC), free fatty acids (FFA), prostaglandin E2 (PGE2) using respective ELISA kits (Meimian), lactate dehydrogenase (Jiancheng Bioengineering Institute, Nanjing, China), as well as insulin, ghrelin, leptin, FGF21, interleukin-6 (IL-6), adrenaline, glucagon, tumour necrosis factor-alpha (TNF•), and C reactive protein (CRP) using respective ELISA kits (Ruixin).

### 2.5. Intraperitoneally glucose and insulin tolerance tests

For the glucose tolerance test, each mouse fasted overnight and was intraperitoneally injected with D-glucose (2 g/kg body weight). For the insulin tolerance test, each mouse fasted for 4 hours and was intraperitoneally injected with insulin (0.5 IU/kg body weight; Beyotime Biotech Inc, Shanghai, China). Blood glucose levels were measured before glucose or insulin injection as 0 min from the tail vein and at 15, 30, 60, 90, and 120 minutes post.

### 2.6. Hepatic parameters measurements

Hepatic tissues (0.1 g) were homogenised and resuspended in PBS. The concentrations of TC and TGs were measured using the corresponding detection kits (Meimian Industrial Co. Ltd., China).

### 2.7. Histological analysis

Histological analysis and estimation of the cellular cross-sectional area followed the standard methods described in our previous study of Pai male mice ^3^.

### 2.8. Western blot

Western blot was performed according to the standard procedures described in our previous study ^3^. Briefly, 20 µg total protein extracts were separated by 10-15% SDS-PAGE gel and transferred to the PVDF membranes (Millipore ISEQ00010, Merck). Each membrane was blocked and cut into 2-3 parts, and then each portion was hybridised with an antibody to the protein of interest or beta-ACTIN (Proteintech Group, Inc, China, 20536-1-AP) or GAPDH (Abcam AB8245), followed by HPR-labelled goat anti-rabbit IgG (Santa Cruz Biotechnology, sc-2004). The chemiluminescent signal was developed using the SuperSignal™ West Femto substrate (ThermoFisher, USA). Blots were imaged for 5 s to 2 min and quantified using ImageJ software (NIH), and values were respectively normalised to beta-ACTIN or GAPDH as a loading control. The primary antibodies were rabbit antibodies targeting AMP-activated protein kinase (AMPK; Abcam, ab207442), adipose triglyceride lipase (ATGL; Proteintech, 55190-1-AP), carnitine palmitoyltransferase I-a (CPT1A; Proteintech, 15184-1-AP) or I-b (CPT1B; Proteintech, 22170-1-AP), fatty acid synthase (FASN; Abcam, ab128870), FGF21 (Abcam, ab171941), phospho-AMPK (pAMPK; Abcam, ab133448), phospho-hormone-sensitive lipase (pHSL; Cell Signaling Technology, #4139), UCP1 (Abcam, ab234430), or UCP2 (Proteintech, 11081-1-AP).

### 2.9. Metabolic cage measurements

Indirect calorimetry, heat production, and activities of mice at 11∼12 weeks of age were measured using the Automated Home Cage Phenotyping TSE PhenoMaster V4.5.3 system (TSE Systems Inc., Germany). Mice were individually housed in plexiglass cages and fed ad libitum. Food intake, VO_2_, and VCO_2_ were measured every 39 minutes. The climate chamber was set to 22°C with a 12-hour light-dark cycle (lights on at 0700 h). After a 48-hour acclimation period, data were collected for the following 72 hours, and the calculated lean mass was adjusted for all measurements.

### 2.10. Statistical analysis

Analysis was conducted using GraphPad Prism 8.0 (GraphPad Software Inc.; La Jolla, CA, USA). Comparisons between wt and Pai/Pai samples were performed using an unpaired two-tailed t-test with Welch’s correction where appropriate. Multiple comparisons among wt, Pai/wt, and Pai/Pai groups were performed using one-way ANOVA and Brown-Forsythe and Welch tests. Data are presented as mean ± standard deviation (SD), and p < 0.05 was considered statistically significant.

## 3. RESULTS

### 3.1. A shift of t10c12-CLA and changes in fatty acids compositions in Pai mice

Real-time analysis revealed that the foreign Pai gene transcribed in WAT, BAT, livers, and hypothamalus of Pai/wt and Pai/Pai mice in a dose-dependent manner (Fig. 1a). Gas chromatography analysis of FA compositions in the livers, kidneys, hearts, tibialis anterior muscle, and interscapular BAT tissues revealed that the content of t10c12-CLA had increased by 69% in the Pai/wt kidneys compared to wt littermates (p < 0.05; Fig. 1b). However, the quantities of substrate linoleic acid had decreased in the Pai/Pai livers by 23% and skeletal muscle by 34% and had increased in the Pai/wt kidneys by 36% (p < 0.05; Fig. 1b). The contents of other FAs, such as myristic (14:0), palmitic (16:0), stearic (18:0), palmitoleic (16:1n-7), cis-vaccenic (18:1n-7), oleic (18:1n-9), arachidonic (20:4n-6), and docosahexaenoic (22:6n-3) acids were altered to varying degrees in one or more tissues (Supplementary Table S3). Additionally, the content of total FAs (mg/g) had increased by 30% in the Pai/wt kidneys (p < 0.05; Fig. 1b), similar to the Pai/wt kidney of male mice ^3^. The results suggest that the t10c12-CLA-induced changes in the content of each FA were genotype- or tissue-specific.

**Fig 1.**
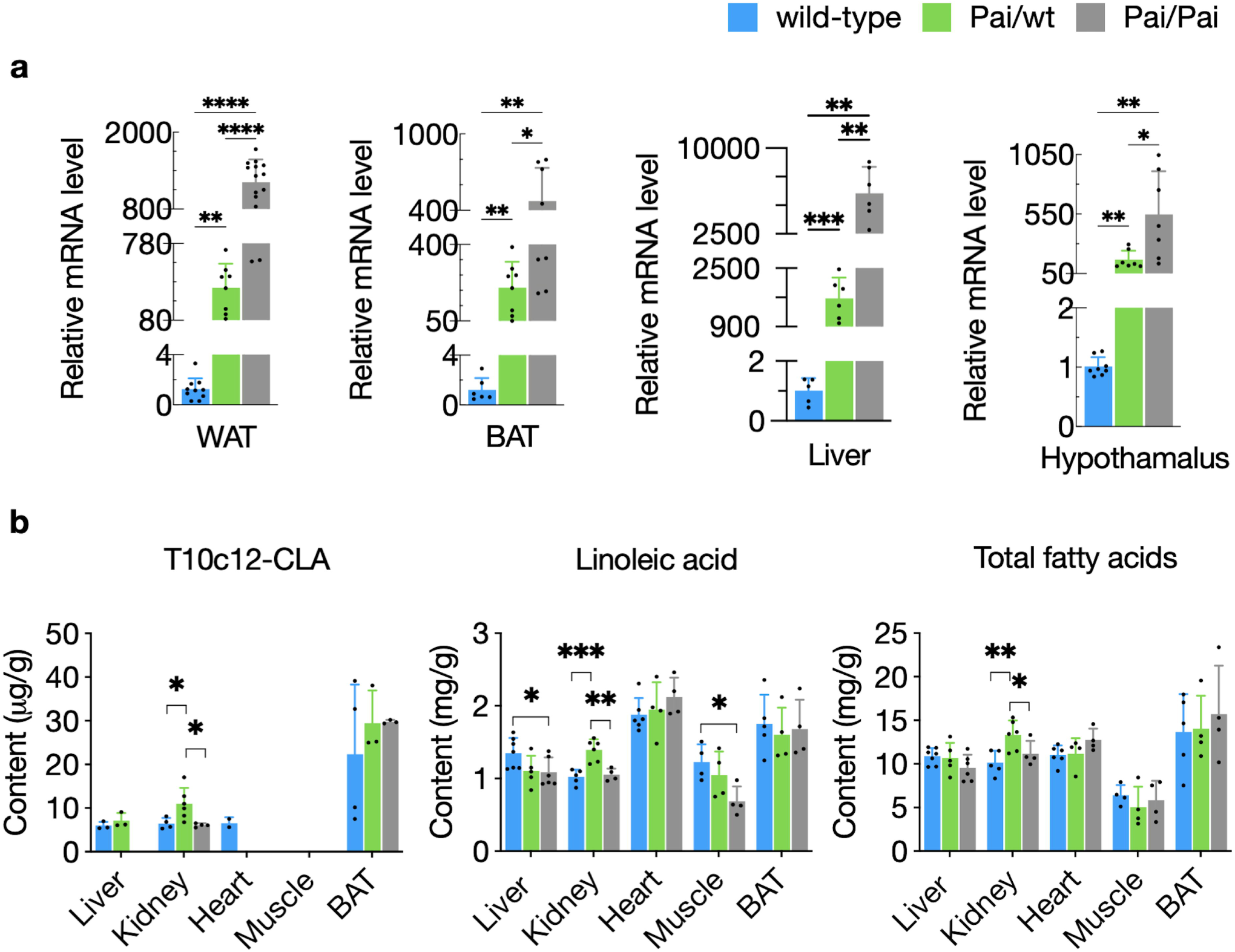
Different mRNA levels of Pai gene (a) and contents of fatty acids (b) in the tissues from wild-type (wt) and Pai mice at the age of 11 weeks. WAT, white adipose tissues; BAT, brown adipose tissue. All fatty acid contents are listed in Supplementary Table S3. The bars represent the mean ± SD. * indicates p < 0.05; ** indicates p < 0.01; *** indicates p < 0.001; and **** indicates p < 0.0001, respectively.

### 3.2. No fat reduction in Pai mice

We first concentrated on the effect of t10c12-CLA on bodyweight and adiposity in Pai mice. The results showed that there was a gradual reduction of weaning weight in Pai/wt and Pai/Pai genotypes compared to wt mice (p < 0.05, Fig. 2a); however, the difference in bodyweight disappeared after five weeks of age and did not appear during the onward ages in both Pai mice (Supplementary Figure S1), similar to the weaning weight of Pai male mice in our previous study ^3^. Magnetic resonance imaging, dissection and histological analyses revealed that Pai/Pai mice at 11 weeks had no reduction of WAT mass and the cross-sectional area of white adipocytes (Fig. 2b-e) but exhibited generalised organomegaly, including significantly enlarged livers, spleens, hearts, and ovaries when body weight was considered (p < 0.05; Supplementary Table S4). Pai/wt mice also had no mass loss of WAT and only showed enlarged ovaries.

**Fig 2.**
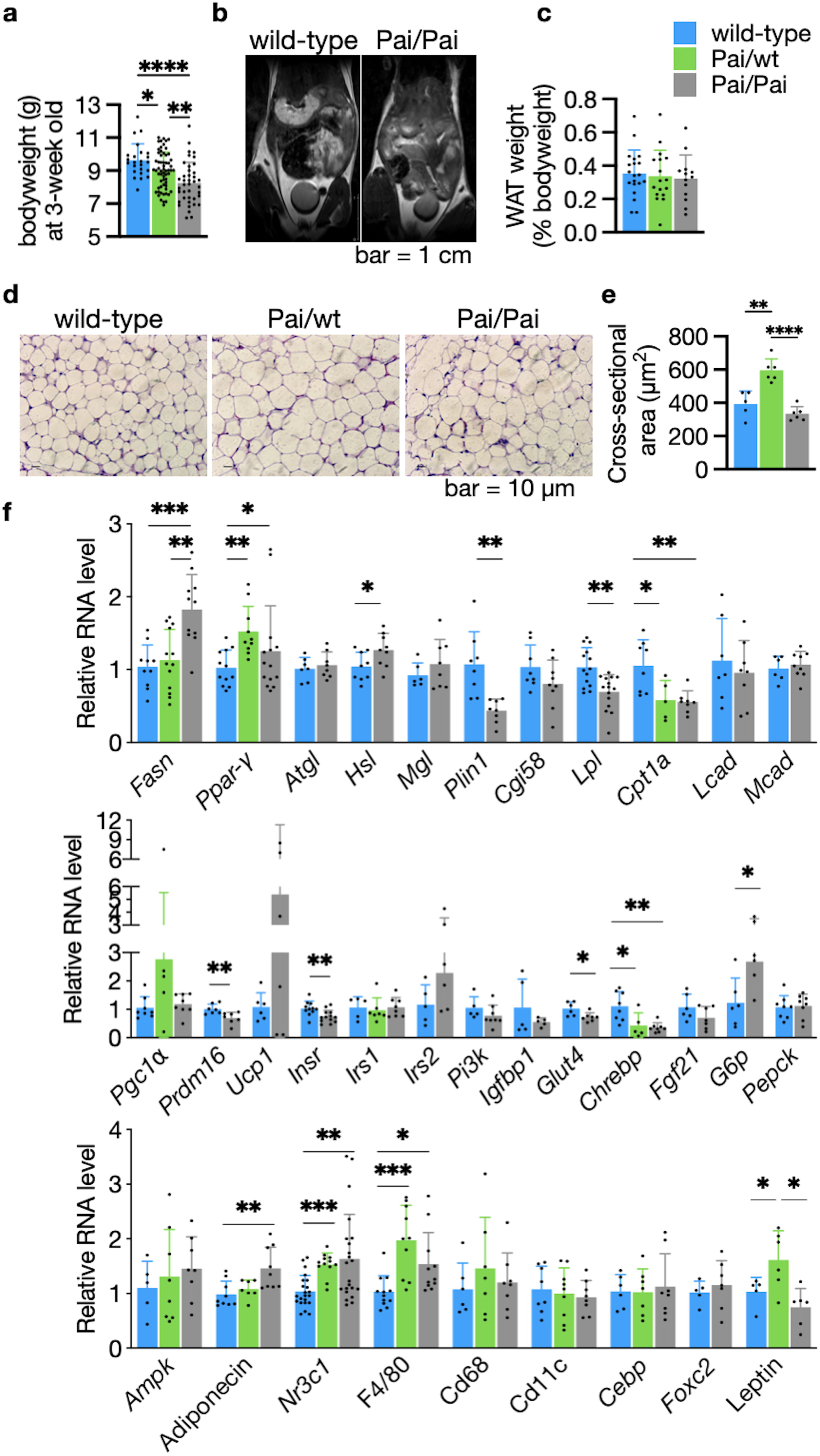
Magnetic resonance imaging and histological and RNA expression analysis of white adipose tissues (WAT). (a) Decrease of weaning weight in Pai mice. (b) Coronal sections were obtained on the whole-body midsection for four wild-type (wt) or Pai/Pai mice at 11 weeks in each group during magnetic resonance imaging. The wt and Pai/Pai mice show similar areas of signal intensity of fatty tissue in the subcutaneous and intra-abdominal regions. (c) There is no difference in the mass of WAT among wt, Pai/wt, and Pai/Pai mice. (d) Imaging of hematoxylin-eosin staining, (e) Analysis of cross-sectional area per cell, and (f) Relative expression of mRNA in the WAT. Ampk, AMP-activated protein kinase; Atgl, Adipose triglyceride lipase; Cd, cluster of differentiation; Cebp, CCAAT/enhancer-binding-protein beta; Cgi58, Comparative gene identification 58; Chrebp, Carbohydrate response element binding protein; Cpt1a, Carnitine palmitoyltransferase I-a; Fasn, Fatty acid synthase; Fgf21, Fibroblast growth factor 21; Foxc2, Forkhead box protein c2; G6p, Glucose-6-phosphatase; Glut4, Glucose transporter type 4; Hsl, Hormone-sensitive lipase; Igfbp1, Insulin-like growth factor binding protein 1; Insr, Insulin receptor; Irs1, Insulin receptor substrate-1; Irs2, Insulin receptor substrate-2; Lcad, Long-chain acyl-CoA dehydrogenase; Lpl, Lipoprotein lipase; Mcad, Medium-chain acyl-CoA dehydrogenase; Mgl, Monoacylglycerol lipase; Nr3c1, Nuclear receptor subfamily 3 group C member 1; Pai, Propionibacterium acnes isomerase; Pepck, Phosphoenolpyruvate carboxykinase; Pgc1IZ, Ppar-gamma coactivator 1 alpha; Pi3k, Phosphoinositide 3-kinase; Plin1, Perilipin 1a; Ppar-•, Peroxisome proliferator-activated receptor-•; Prdm16, PR domain containing 16; Ucp1, Uncoupling protein 1. The bars represent the mean ± SD. * indicates p < 0.05; ** indicates p < 0.01; *** indicates p < 0.001; and **** indicates p < 0.0001, respectively.

To investigate in-depth changes in WAT, RNA levels of 33 essential genes were determined in Pai/Pai mice and the total RNA levels of 14 (42%) genes, including seven up-regulated and seven down-regulated, were modified among them (Fig. 2f). Briefly, the seven up-regulated genes were Fasn, Peroxisome proliferator-activated receptor (Ppar)-•, Hsl, glucose-6-phosphatase (G6p), F4/80, Adiponectin, and nuclear receptor subfamily 3 group C member 1 (Nr3c1, which is the corticosterone receptor). The seven down-regulated genes were lipoprotein lipase (Lpl), Cpt1a, perilipin 1a (Plin1), PR domain containing 16 (Prdm16), insulin receptor (Insr), glucose transporter type 4 (Glut4), and carbohydrate response element binding protein (Chrebp). In Pai/wt adipocytes, the RNA levels of six (43%) genes were changed among 14 examined genes. There were up-regulated Ppar-•, Nr3c1, F4/80, and Leptin, and down-regulated Cpt1a and Chrebp (Fig 2f). These aberrant gene expressions in white adipocytes suggest that t10c12-CLA can affect a series of metabolic processes, such as lipid metabolism, lipid and glucose uptake, energy expenditure, stimuli response and glucose homeostasis.

### 3.3. Enhanced lipolysis and thermogenesis in BAT of Pai mice

To clarify the effect of t10c12-CLA on BAT thermogenesis, we measured the BAT features of Pai mice. Magnetic resonance imaging and dissection analysis did not reveal any mass changes of BAT in both Pai mice (Fig. 3a-b). Hematoxylin-eosin staining showed Pai/Pai adipocytes with uniform and small-sized lipid droplets and Pai/wt adipocytes with irregular-sized lipid droplets compared to wt adipocytes with uniform and large-sized lipid droplets (Fig. 3c). The reduced cross-sectional areas per adipocyte indicated that the volumes of brown adipocytes decreased significantly in both Pai genotypes (p < 0.05; Fig. 3d).

**Fig 3.**
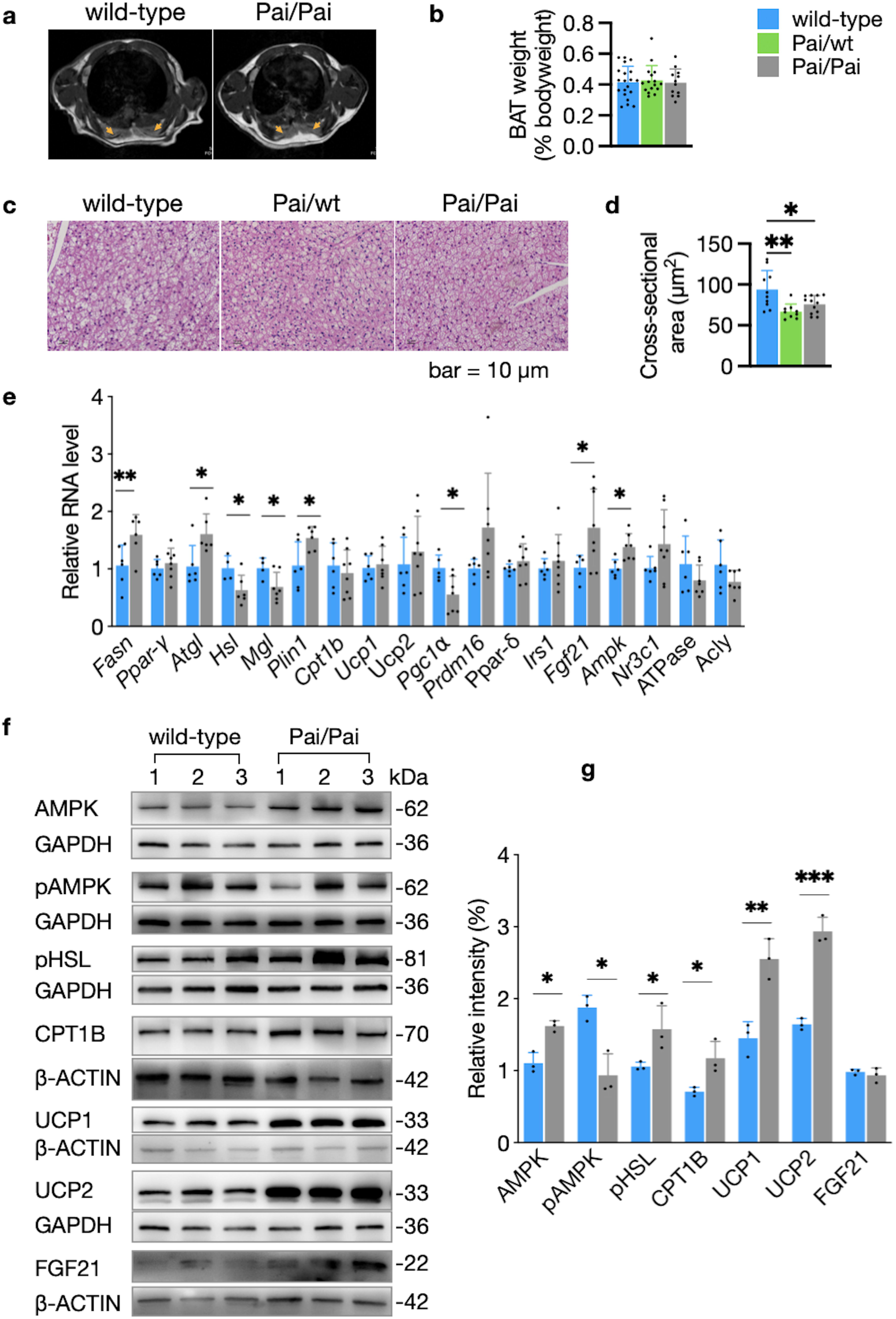
Aspects of brown adipose tissues (BAT). (a) Axial sections of the interscapular BAT show similar areas of grey signal intensity on magnetic resonance images (brown arrows) between four wild-type (wt) and four Pai/Pai mice at 11 weeks. (b) There is no BAT mass difference among wt, Pai/wt, and Pai/Pai mice. (c-d) Analysis of hematoxylin-eosin staining shows that the sizes of lipid droplets become small in the Pai/wt or Pai/Pai adipocytes, and the cross-sectional area per cell is decreased in Pai/wt and Pai/Pai mice. (e) Analyses of the relative expression of mRNAs of critical genes in wt and Pai/Pai mice. (f-g) Western blot analysis of proteins and their relative intensities in wt and Pai/Pai mice. Each membrane was cut into 2-3 parts, and each portion was then hybridised with a corresponding antibody in the western blot. All original, replicated blots were provided in the Supplementary information file. Acly, ATP citrate lyase; Ampk, AMP-activated protein kinase; Atgl, Adipose triglyceride lipase; Cpt1b, Carnitine palmitoyltransferase I-b; Fasn, Fatty acid synthase; Fgf21, Fibroblast growth factor 21; Irs1, Insulin receptor substrate-1; Mgl, Monoacylglycerol lipase; Nr3c1, Nuclear receptor subfamily 3 group C member 1; pAMPK, phosphorylated AMPK; Pgc1IZ, Ppar-gamma coactivator 1 alpha; pHSL, phosphorylated hormone-sensitive lipase; Plin1, Perilipin 1a; Ppar-• and -δ, Peroxisome proliferator-activated receptor-• and -δ; Prdm16, PR domain containing 16; Ucp1 and 2, Uncoupling protein 1 and 2. The bars represent the mean ± SD. * indicates p < 0.05; ** indicates p < 0.01; and *** indicates p < 0.001, respectively.

RNA analysis of Pai/Pai BAT showed that the transcriptional levels of eight genes were changed among 18 tested genes, consisting of up-regulated Fasn, Atgl, Plin1, Fgf21, and Ampk, as well as down-regulated Hsl, monoacylglycerol lipase (Mgl), and ppar-• coactivator 1 alpha (Pgc1•) (p < 0.05; Fig. 3e). In contrast, the other critical genes involved in lipolysis and thermogenesis, such as Ppar-•, Ucp1, Ucp2, Prdm16, and Ppar-•, remained unchanged RNA levels. Western blot analysis revealed over-expressed AMPK, pHSL, CPT1B, UCP1, UCP2, and as well as down-expressed pAMPK in Pai/Pai BAT (p < 0.05; Fig. 3f-g). These results indicate that the t10c12-CLA can stimulate lipolysis, beta-oxidation, and thermogenesis of BAT.

### 3.4. Hepatic steatosis in Pai/Pai mice

Whether t10c12-CLA leads to hepatic steatosis is a conflict in the previous studies. In the current study, Pai/Pai mice exhibited hepatic hypertrophy, unchanged TC concentrations, and increased TGs levels in the livers (p < 0.05; Fig. 4a-c). Histological staining showed profound fat accumulation (∼2.3 fold) in swollen hepatocytes (p < 0.05; Fig. 4d-g), indicating hepatic steatosis in Pai/Pai mice. However, the above parameters of Pai/wt livers remained unchanged compared to wt samples. The results suggest that hepatic steatosis is associated with high doses of t10c12-CLA.

**Fig 4.**
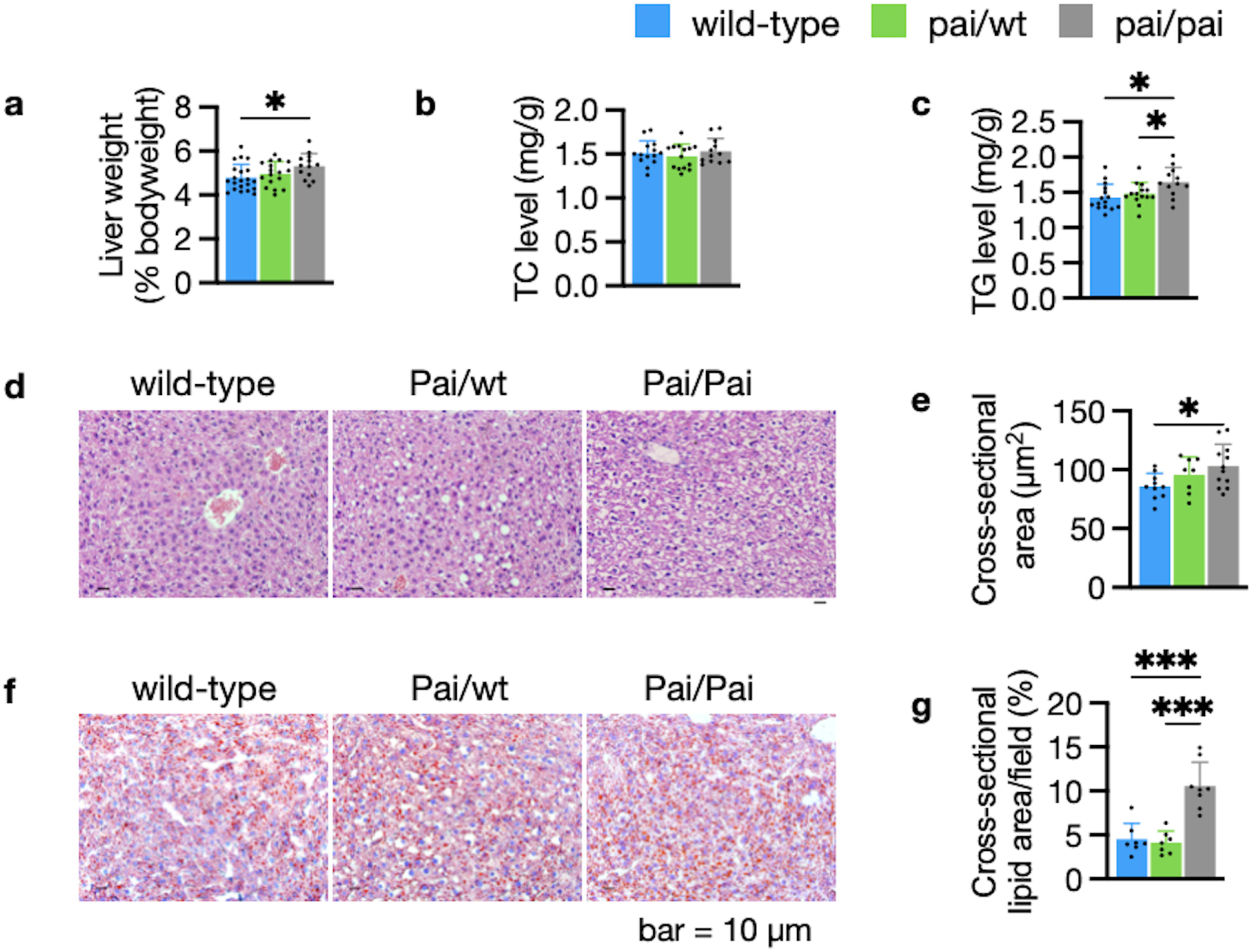
Histological and lipid analysis of livers. (a) Dissection analysis shows an increased liver mass in Pai/Pai mice. (b-c) ELISA analyses indicate normal levels of hepatic total cholesterol (TC) and increased levels of triglycerides (TGs) in Pai/Pai livers. (d-e) Analyses of hematoxylin-eosin staining show the abnormal morphology and hepatocyte oedema and the enlarged cross-sectional area per hepatocyte in Pai/Pai mice. (f-g) Analyses of oil red staining show hepatic lipid accumulation in Pai/Pai mice. The bars represent the mean ± SD. * indicates p < 0.05 and *** indicates p < 0.001, respectively.

RNA analysis of Pai/Pai livers revealed that 25 (50%; Fig. 5a) genes significantly (p < 0.05) altered their RNA levels among 50 tested genes, Briefly, genes involved in lipid metabolism included three up-regulated Fasn, Mgl, and comparative gene identification 58 (Cgi58) and eight down-regulated genes Ppar-•, diacylglycerol acyltransferase (Dgat) 1, Dgat2, Atgl, Lpl, Cpt1a, medium-chain acyl-CoA dehydrogenase (Mcad), and acyl-CoA oxidase (Acox1). Genes involved in the sterol pathway included up-regulated sterol regulatory element binding protein (Srebp) 1c and insulin-induced gene (Insig) 1, as well as down-regulated Srebp1a, Insig2, HMG-CoA reductase (Hmgcr), and LDL receptor (Ldlr). Moreover, genes related to insulin/insulin-like growth factor (IGF) signalling and glucose metabolism included up-regulated forkhead box protein a2 (Foxa2), as well as down-regulated IGF bind protein 1 (Igfbp1), Glut4, Fgf21, G6p, and phosphoenolpyruvate carboxykinase (Pepck). Additionally, two NADPH-producing enzymes, Malic and 6-Phosphogluconate dehydrogenase (Pgd), were up-regulated, while the inflammatory factor cluster of differentiation 11c (Cd11c) was down-regulated in the Pai/Pai livers. Western blot analysis revealed that the protein levels of pAMPK and FASN were reduced, and CPT1A was increased, while AMPK, ATGL, CPA1B, and FGF21 levels remained unchanged (Fig. 5b-c).

**Fig 5.**
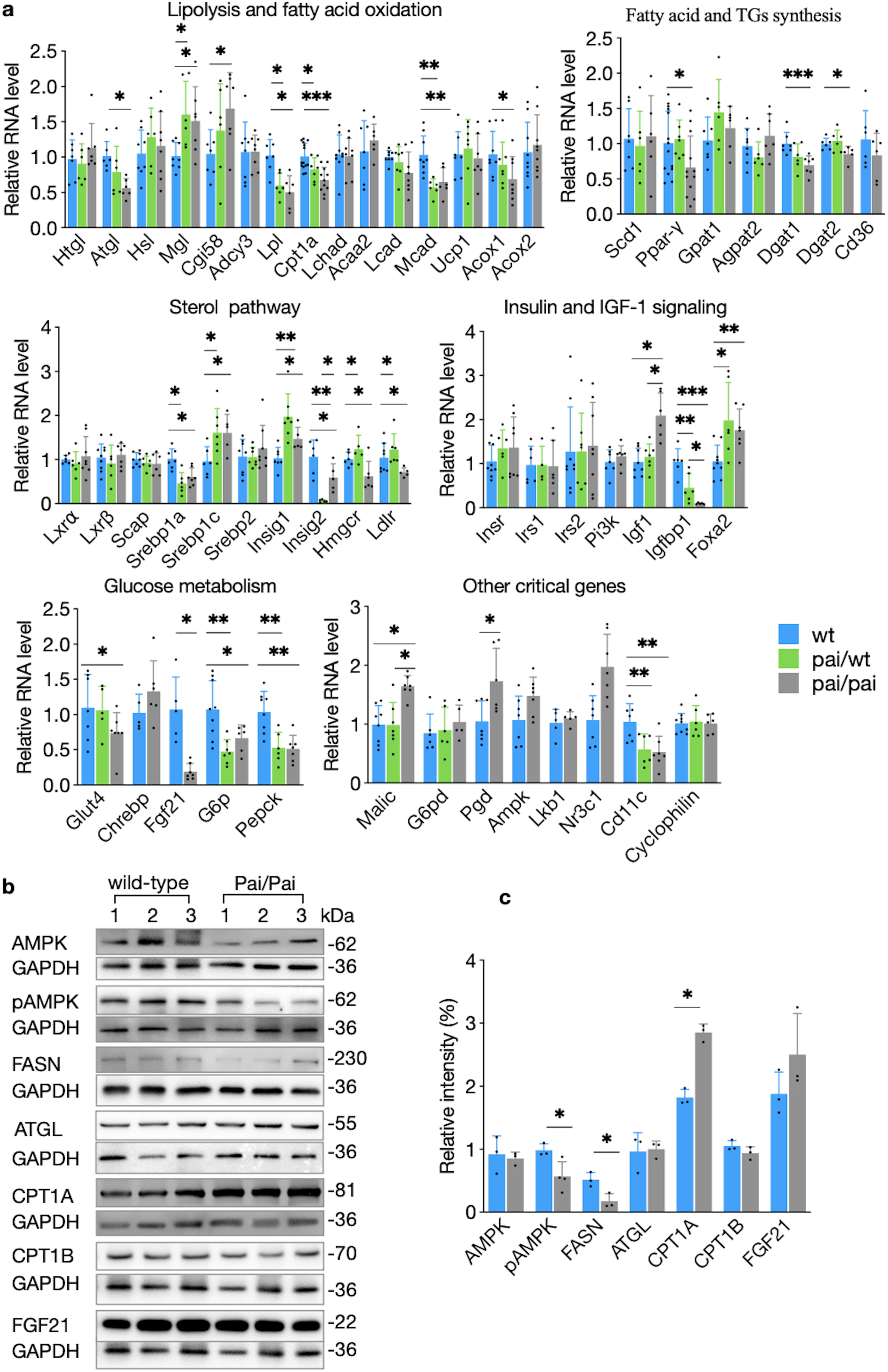
Analysis of gene expression in the livers. (a) Analyses of the relative expression of mRNAs of critical genes in wild-type, Pai/wt, and Pai/Pai mice. (b-c) Western blot analysis of proteins and their relative intensities in wild-type and Pai/Pai mice. Each membrane was cut into 2-3 parts, and each portion was then hybridised with a corresponding antibody in the western blot. All original, replicated blots were provided in the Supplementary information file. Acca2, Acetyl-CoA acyltransferase 2; Acox, Acyl-CoA oxidase; Adcy3, Adenylate cyclase 3; Agpat2,; 1-acylglycerol-3-phosphate O-acyltransferase 2; Ampk, AMP-activated protein kinase; Atgl, Adipose triglyceride lipase; Cd, cluster of differentiation; Cebp, CCAAT/enhancer-binding-protein beta; Cgi58, Comparative gene identification 58; Chrebp, Carbohydrate response element binding protein; Cpt1a, Carnitine palmitoyltransferase I-a; Dgat, Diacylglycerol acyltransferase; Fasn, Fatty acid synthase; Fgf21, Fibroblast growth factor 21; Foxa2, Forkhead box protein a2; G6p, Glucose-6-phosphatase; G6pd, Glucose-6-phosphate dehydrogenase; GAPDH, glyceraldehyde-3-phosphate dehydrogenase; Glut4, Glucose transporter type 4; Gpat1, Glycerol-3-phosphate acyltransferase 1; Hmgcr, HMG-CoA reductase; Hsl, Hormone-sensitive lipase; Htgl, Hepatic triglyceride lipase; Igf1, Insulin-like growth factor-1; Igfbp1, IGF binding protein 1; Insig 1 and 2, Insulin induced gene 1 and 2a; Insr, Insulin receptor; Irs, Insulin receptor substrate; Lcad, Long-chain acyl-CoA dehydrogenase; Lchad, Long-chain 3-hydroxyacyl-CoA dehydrogenase; Ldlr, LDL receptor; Lkb1, liver kinase B1; Lpl, Lipoprotein lipase; Lxr, Liver X receptor; Mcad, Medium-chain acyl-CoA dehydrogenase; Mgl, Monoacylglycerol lipase; Nr3c1, Nuclear receptor subfamily 3 group C member 1; Pai, Propionibacterium acnes isomerase; Pepck, Phosphoenolpyruvate carboxykinase; Pgc1IZ, Ppar-gamma coactivator 1 alpha; Pgd, 6-Phosphogluconate dehydrogenase; Pi3k, Phosphoinositide 3-kinase; Plin1, Perilipin 1a; Ppar-•, Peroxisome proliferator-activated receptor-•; Prdm16, PR domain containing 16; Scap, SREBP cleabage-activating protein; Srebp, Sterol regulatory element binding protein; Scd1, Stearoyl-CoA desaturase 1; Ucp1, Uncoupling protein 1. The bars represent the mean ± SD; * indicates p < 0.05; ** indicates p < 0.01; and *** indicates p < 0.001, respectively.

RNA analysis of Pai/wt livers showed that 32 genes had similar transcription levels to Pai/Pai samples, and 13 (33%) of them were markedly (p < 0.05) different from those in wt samples among 40 tested genes (Fig. 5a). To be specific, these 13 genes included four up-regulated (Mgl, Srebp1c, Insig1, and Foxa2) and nine down-regulated (Lpl, Cpt1a, Mcad, Srebp1a, Insig2, Igfbp1, G6p, Pepck, and Cd11c). Furthermore, RNA results indicated that the mRNA levels of almost all genes in the Pai/wt sample were intermediate between those of wild-type and Pai/Pai mice, and the RNA levels of most genes decreased in Pai’s liver, suggesting that the 10c12-CLA attenuates glucose and lipid metabolism in a dose-dependent manner.

### 3.5. Changes in hormones, triglycerides and inflammatory factors in Pai mice

Although the circulating levels of TC, FFAs, and HDL showed no differences between Pai mice and their wt littermates, the levels of plasma TGs were significantly elevated in Pai/Pai mice compared to wt or Pai/wt mice (p < 0.05, Fig. 6), indicating t10c12-CLA-induced hypertriglyceridemia in Pai/Pai mice. However, the concentrations of circulating glucose, insulin, leptin, and ghrelin (Fig. 6), as well as small intestine length (Supplementary Table S4), showed no differences (p > 0.05) between Pai mice and their wt littermates, suggesting no effect of t10c12-CLA on energy intake.

**Fig 6.**
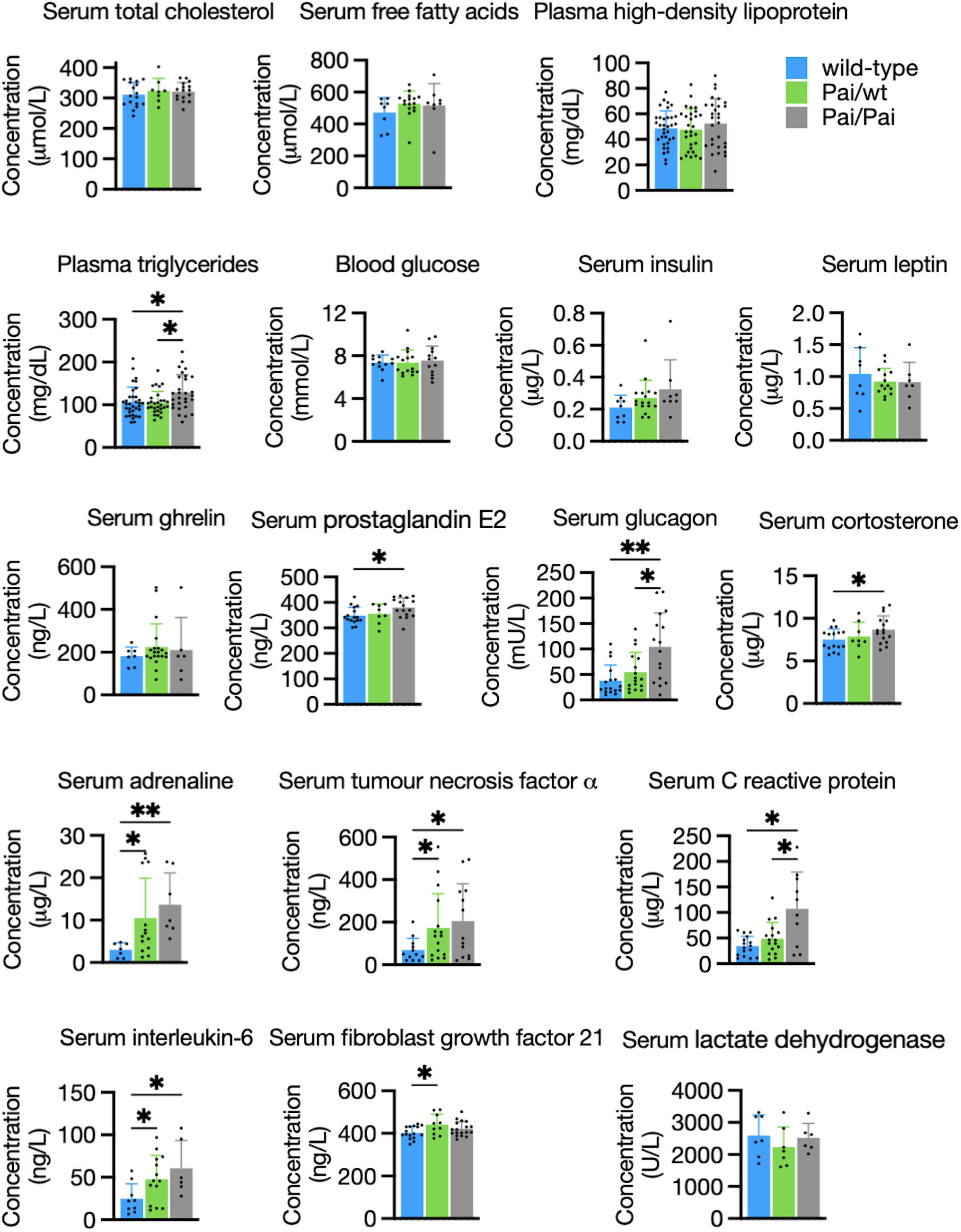
Comparisons of circulating factors in wild-type and Pai mice. Blood samples were collected from non-fasted mice at the age of 11∼15 weeks. The bars represent the mean ± SD. * indicates p < 0.05; ** indicates p < 0.01; and *** indicates p < 0.001, respectively.

Some critical hormones related to lipid metabolism and inflammation were also investigated. Serum concentrations of PGE2 (109%), glucagon (2-fold), corticosterone (116%), adrenaline (4.5-fold), TNF• (3-fold), CRP (3-fold), and IL-6 (2.5-fold) were significantly (p < 0.05) increased in Pai/Pai mice when compared to their wt littermates. However, only adrenaline (3.5-fold), TNF• (2.5-fold), and IL-6 (2-fold) were significantly (p < 0.05) elevated in Pai/wt mice. Additionally, the levels of all the above factors in Pai/wt mice were intermediate between those of wild-type and Pai/Pai mice, suggesting that the effect of 10c12-CLA on hormone overproduction is dose-dependent. The only exception is FGF21, which was exclusively elevated by 9% in Pai/wt, not in Pai/Pai female mice (Fig. 6), similar to the previous result in male mice ^3^, suggesting that FGF21 is sensitive to low-dose, not high-dose t10c12-CLA. Moreover, there was no increase in blood levels of lactic acid dehydrogenase in either Pai mice, indicating no cellular injury caused by the PAI protein.

Both glucose and insulin tolerance tests showed no difference in the circulating glucose concentrations or the area under the curve between wt and two Pai genotypes (Fig. 7a-b). Interestingly, the fasted glucose levels before glucose injection were significantly (p < 0.05) higher in Pai/wt mice than in wt or Pai/Pai mice (Fig. 7a). So we further measured blood glucose levels during a 24-hour fasting duration to investigate this discrepancy. We found that Pai/wt mice continuously maintained higher glucose levels than their wt littermates (p < 0.05; Fig. 7c). This suggests that glucose stability in starved Pai/wt mice is sensitive to low doses of t10c12-CLA, which may be associated with the beneficial insulin-sensitising and glucose-lowering effects of FGF21 ^19^.

**Fig 7.**
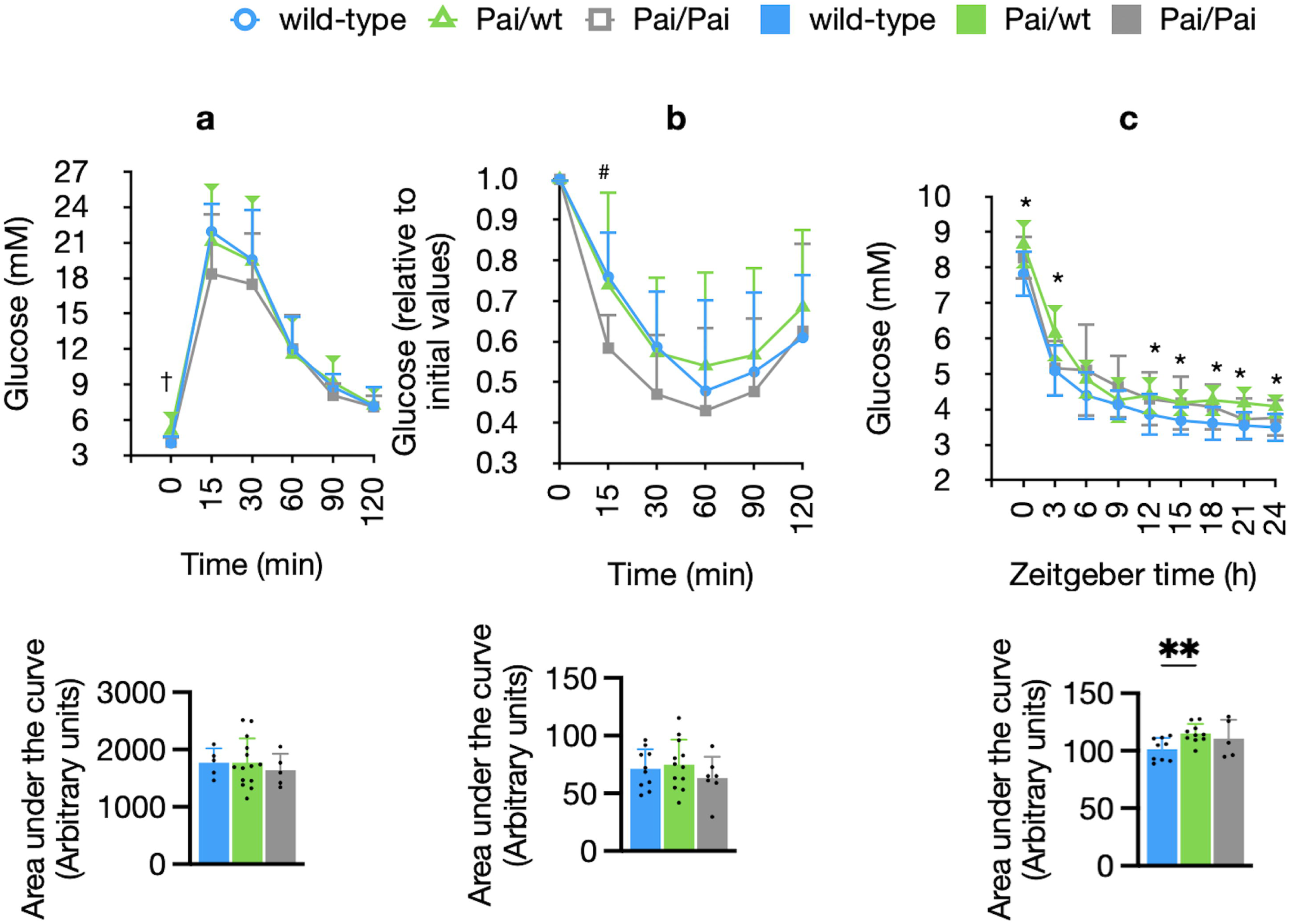
Comparisons of glucose (a) and insulin (b) tolerance tests and dynamic glucose levels (c) in wild-type (wt) and Pai mice. (a) Absolute blood glucose levels and the area under the curve. (b) Blood glucose levels relative to initial values and the area under the curve. (c) Dynamic blood glucose levels and the area under the curve are measured in fasting mice starved from Zeitgeber time 0 to 24. Zeitgeber times 0 and 12 are lights-on and -off times, respectively. The bars represent the mean ± SD. † indicates p < 0.05 between Pai/wt and wt or Pai/Pai mice; # indicates p < 0.05 between wt and Pai/Pai mice; * indicates p < 0.05 between wt and Pai/wt mice.

### 3.6. Less heat release, oxygen consumption, and physical activities in Pai mice

To investigate the potential impact of t10c12-CLA on energy homeostasis, energy metabolism and activity parameters were assessed using metabolic cages, considering body weight. Compared to their wt littermates, both Pai/wt and Pai/Pai mice had no differences in body weight gain, food and water intake (Fig. 8a-c), total distance travelled during the whole 72-h observation (Fig. 8d), and respiratory exchange ratio (Fig. 8e-f). However, the following parameters showed varying differences.

**Fig 8.**
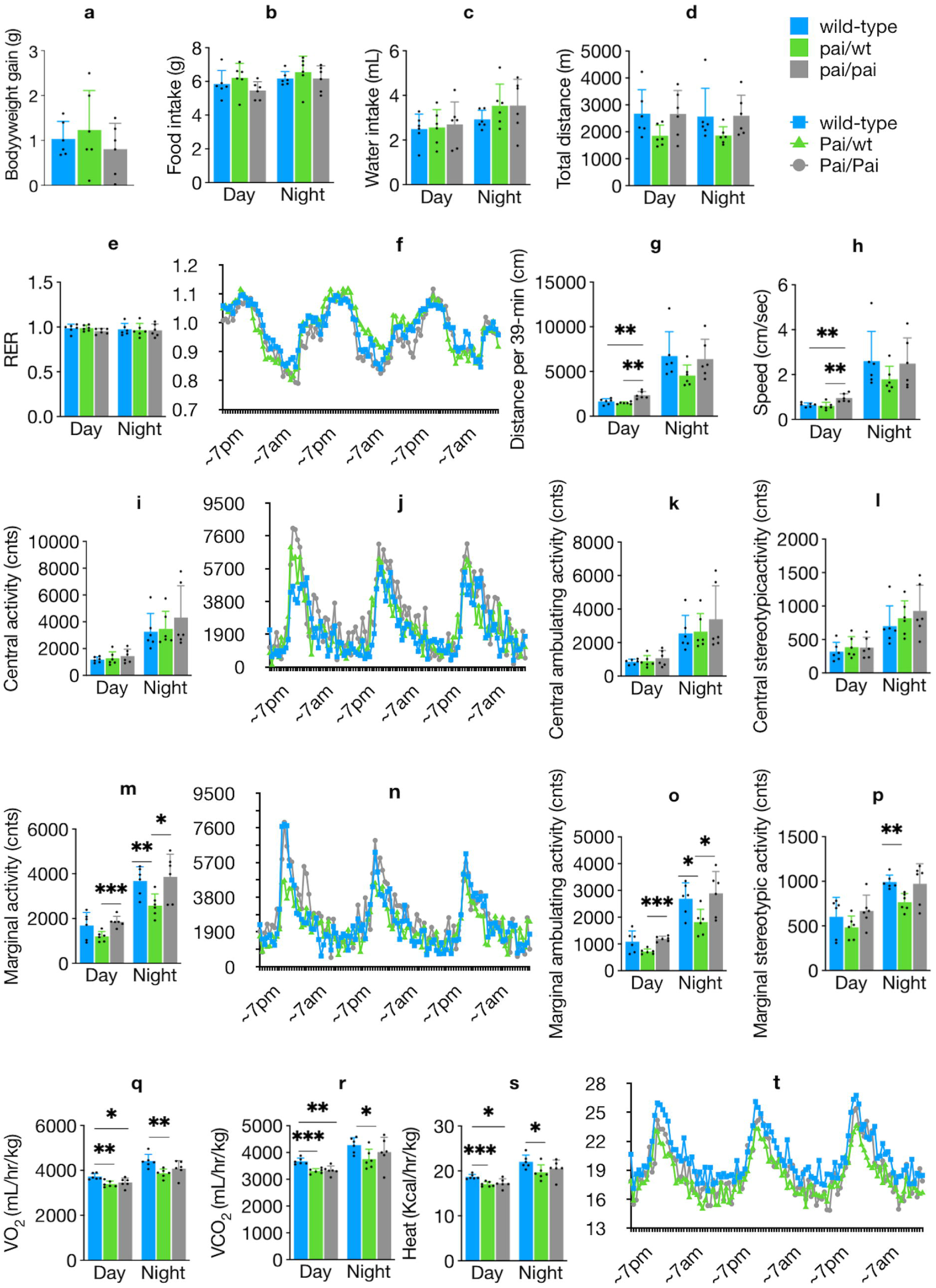
Abnormal energy metabolism and activities in Pai mice. Comparison of body weight gain during a 72-h period (a), food intake (b), water intake (c), total distance travelled during a 72-h period (d), respiratory exchange ratio (RER) and RER time course (e-f), distance travelled during a 39-min sampling duration (g), speed (h), locomotor activities in the centre of the cage and its time course (i-j), central ambulating and stereotypic activities (k-l), locomotor activities in the margin or corners of the cage and its time course (m-n), marginal ambulating and stereotypic activities (o-p), O_2_ consumption (q), CO_2_ production (r), heat release and its time course (s-t) among wild-type, Pai/wt, and Pai/Pai mice at 11 weeks. The lights turned on and off at 7:00 am and 7:00 pm. The data are normalised to lean body weight. The bars represent the mean ± SD. * indicates p < 0.05; ** indicates p < 0.01; and *** indicates p < 0.001, respectively.

The Pai/Pai mice covered a longer distance per sampling duration (∼44% longer) and exhibited 47.2% higher speed during the light phase (p < 0.05; Fig. 8g-h) but did not reduce their total locomotor activities in either the centre (Fig. 8i-l) or margin/corners (Fig. 8m-p). Nevertheless, during both the light (p < 0.05) and dark (p > 0.05) periods, they consumed 7.1% and 7.6% less O_2_ (Fig. 8q) and produced 9.3% and 5.9% less CO_2_ (Fig. 8r) as well as 7.6% and 6.9% less heat (Fig. 8s-t), respectively. The results suggest that Pai/Pai mice simultaneously reduced oxygen consumption and heat release during the light periods while maintaining normal locomotor activities.

On the other hand, Pai/wt females did not reduce their central activities (p > 0.05) but displayed 29.9% fewer marginal activities, such as marginal ambulating and stereotypic activities during the dark period (p < 0.05). Moreover, during both the light (p < 0.05) and dark (p < 0.05) periods, they consumed 9.0% and 12.4% less O_2_, produced 9.9% and 12.0% less CO_2_, as well as 8.9% and 10.7% less heat, respectively, suggesting that Pai/wt mice simultaneously reduced oxygen consumption and heat release during the whole day while exhibiting fewer marginal activities.

### 3.7. Abnormal gene transcription in the hypothalamus of Pai mice

To determine whether t10c12-CLA affected the hypothalamus, critical hypothalamic genes and proteins were also measured. The mRNA levels of the leptin receptor and agouti-related peptide increased, and the transcription levels of the ghrelin receptor and Orexins decreased in the Pai/wt or Pai/Pai hypothalamus compared to wt mice (Supplementary Figure S2). Western blot analysis had not revealed any changes in AMPK, phosphorylated AMPK, and glucose-related protein 78 (GRP78) in the Pai/Pai hypothalamus; and the ratio of pAMPK/AMPK had no difference between wt and Pai/Pai samples (p > 0.05; Supplementary Figure S2). These findings suggest that the central regulation of energy intake may only be affected at the transcriptional level.

## 4. DISCUSSION

Our previous observation showed that t10c12-CLA reduced body fat by releasing heat via BAT activation with lower blood TGs in Pai/Pai male mice ^3^. Conversely, in the current study, t10c12-CLA did not cause body fat loss but resulted in complicated symptoms in female mice, including a dose-dependent increase in multiple hormones and inflammatory factors (PGE2, glucagon, corticosterone, adrenaline, IL-6, and TNF•), less heat release, fatty liver and hypertriglyceridemia. It suggests there is a sexually different impact of t10c12-CLA on health. However, the excess FGF21 in Pai/wt mice in both sexes suggests that FGF21 secretion is sensitive to low-dose t10c12-CLA.

The metabolic pathway of t10c12-CLA, similar to that of linoleic or alpha-linolenic acids ^20^, suggests it can competitively interrupt the metabolic pathway of n-6 or n-3 FAs and their derived leukotrienes signals/hormonal molecules. It also indicates that t10c12-CLA can be directly converted into the analogues of n-6 FAs and their derivatives. PGE2, an arachidonic acid-derived hormone, will be easily disturbed by t10c12-CLA. PGE2 is involved in stress response and has essential effects on inflammation, fever, and renal filtration ^21^. It can induce corticosterone secretion by regulating the hypothalamic-pituitary-adrenal axis activity and enhancing adrenaline synthesis in the Chromaffin cells of the adrenal medulla. Subsequently, the raised circulating corticosterone and adrenaline could stimulate excess glucagon secretion ^22^. In Pai/Pai female mice, we have observed overproduction of PGE2 induced by t10c12-CLA and inferred that the excessive PGE2 (or analogues) induced the increases in corticosterone, glucagon, adrenaline, and inflammatory factors (IL-6, TNFIZ, and CRP). As a matter of fact, less heat release in Pai/Pai female mice may be related to hyperprostaglandinemia, hyperadrenerginemia and inflammatory response, while hyperglyceridemia and fatty liver are associated with hypercorticosteroidemia and glucagon resistance.

In the hypothalamic preoptic area, PGE2 receptors EP3, coupled to cAMP decreases, can stimulate a passive defence mechanism resembling less O_2_ consumption and less heat release to avoid further energy expenditure. In contrast, receptors EP4, associated with cAMP increases, can stimulate an active defence mechanism, resembling more O_2_ consumption and febrile responses through increasing BAT thermogenesis, which typically occurs to promote well-being and survival ^21,23^. It suggests that gender-based differences in PGE2 levels might partially result in the opposite heat release pattern in male and female Pai/Pai mice. In female mice, the co-elevation of PGE2 and corticosterone suggests that t10c12-CLA induced a passive defence mechanism, including less heat production, which conversely promotes BAT thermogenesis and lipid metabolism via negative feedback to down-regulate pAMPK and up-regulate UCP1, UCP2, CPT1B, and pHSL. In contrast, the lack of PGE2 elevation in male mice suggests that t10c12-CLA induced an active defence mechanism, including hyperthermia, to promote survival ^3^.

In addition to PGE2- and adrenaline-induced heating ^24^, BAT thermogenesis can be mediated through the central GRP78 ^25,26^ or AMPK-OREXINS-BMP8B pathway ^27^. Orexin-null mice exhibited periods of decreased activity during the dark phase and demonstrated less heat production, reduced oxygen consumption, and blunted thermogenic responses ^28^; furthermore, BMP8B’s thermogenic effect is sexually dimorphic and observed in females only ^27^. The Pai/Pai female mice exhibited similar phenotypes to orexin-null mice and lower transcription levels of Orexins in the Pai hypothalamus. These characteristics suggest that the t10c12-CLA-induced BAT thermogenesis may be regulated through central pathways in female mice. Unfortunately, we did not observe evidential changes in GRP78, AMPK, pAMPK, and pAMPK/AMPK ratio in the hypothalamus and could not infer that BAT thermogenesis is associated with the central network controlling energy homeostasis.

Previous studies on mice have yielded conflicting results on whether dietary t10c12-CLA leads to hepatic steatosis ^11,16,29,30^. The current observation of fatty liver and hypertriglyceridemia in Pai/Pai, not in Pai/wt mice, suggests that the t10c12-CLA-induced steatosis is dose-dependent. High-dose t10c12-CLA-induced steatosis may be associated with excess corticosterone because it can lead to Cushing syndrome, corticosteroid-induced lipodystrophy, hypertriglyceridemia, or hepatic steatosis ^31,32^. Another reason for steatosis is the existence of glucagon resistance because it can increase liver fat ^33-35^. The chronic increase in plasma glucagon promotes lipolysis through INSP3R1/CAMKII/ATGL ^36^ or PKA/HSL pathways in hepatocytes ^37^ and stimulates beta-oxidation by activating AMPK/PPAR-IZ/CPT1A/MCAD pathway ^22^. In the current study, the serum glucagon concentration increased nearly twofold in Pai/Pai female mice, similar to the corresponding physiological increase (2∼3 fold) in response to long-term fasting or hypoglycemia ^38^. In contrast, the protein levels of unchanged ATGL, decreased pAMPK, and increased CPT1A indicate that the chronic glucagon excess did not stimulate lipolysis and beta-oxidation in the liver efficiently. These results suggest that the chronic impact of t10c12-CLA might give rise to glucagon resistance in Pai/Pai female mice. In this case, the down-regulated hepatic enzymes may not fully and immediately catalyse the FAs molecules mobilised from other tissues based on glucagon stimulation. Consequently, it disrupts hepatic and systemic lipid homeostasis, leading to dyslipidemia and fatty liver in female mice. Contrarily, the t10c12-CLA did not cause the overproduction of corticosterone or glucagon in Pai/Pai male mice. Thus, the male mice had no fatty liver disease and even exhibited lower TGs levels in the blood.

FGF21 is a vital regulator of the specific hormonal signalling pathway in increasing energy expenditure, reducing hepatic and plasma TGs levels, accelerating lipoprotein catabolism in WAT and BAT ^39^, and suppressing physical activity via central action ^40^. FGF21 signalling enhances insulin sensitivity and lowers serum glucose by acting directly on adipose tissues ^19,41^. The Pai/wt mouse is the first animal in which t10c12-CLA can induce FGF21 secretion, indicating that FGF21 secretion is associated explicitly with low-dose t10c12-CLA in both genders. We do not know if the FGF21 secretion is an early pathological change of hepatic steatosis per se, for liver damage might lead to increased FGF21 release. In turn, if it is a beneficial stimulus from t10c12-CLA, then the health benefits of low-dose t10c12-CLA may be due, in part, to its ability to stimulate FGF21 secretion. Therefore, the different effects of t10c12-CLA on glucose metabolism in diabetic Zucker fatty rats ^42^ and insulin resistance in mice ^12,43^ may be related to the different levels of FGF21 induced by t10c12-CLA. In the current study, excess FGF21 in Pai/wt mice may be associated with specific phenotypes, such as blood glucose homeostasis during starvation. The exact mechanism by which t10c12-CLA activates FGF21 secretion remains unclear and requires further investigation.

Based on the excesses of hormones, abnormality of the energy metabolism, and physical activities in Pai mice in this study, we suggest that t10c12-CLA easily induces excessive adrenaline and results in inflammation reaction, lower energy consumption, and less heat production in lean female mice (Supplementary Figure S3, middle panel). Simultaneously, when high doses of t10c12 are used, it can specifically result in an increased PEG2, which probably induces increases in corticosterone and glucagon, and the combination of these factors causes metabolic syndrome, including fatty liver and hyperglyceridemia in female mice (Supplementary Figure S3, right panel). Conversely, when low doses of t10c12-CLA are used, it can specifically cause an increase in circulating FGF21, which can accelerate lipoprotein catabolism in adipocytes and suppress physical activity (Supplementary Figure S3, left panel). For t10c12-CLA has system-wide effects on multiple tissues via various hormones in female animals, interpreting the current data is challenging and requires careful consideration and more work is needed to understand the metabolic impacts of t10c12-CLA.

Overall, the effect of t10c12-CLA is interfering with lipid metabolism. It results in fat reduction in males but unfavourable metabolic effects in females when fed with a chow diet. More complexly, the metabolism of each tissue may be regulated by multiple hormones through various metabolic pathways. Unfortunately, we cannot judge which tissue or hormone is the primary target of t10c12-CLA and which others are subordinate or passive at this moment. Here, we can only provide as many appearances as possible to provide clues for in-depth research on the impact of t10c12-CLA on female health.

## Acknowledgement

This work was supported by the National Key Research and Development Program of China (2018YFC1003800) and the Priority Academic Program Development of Jiangsu Higher Education Institutions (PAPD). YR received an award from the Jiangsu Funding Program for Excellent Postdoctoral Talent. We thank graduate students Qi Wang, Mei Liu, and Jiaqi Lu in our lab for their generous assistance in partial tests and Ms Yuyang Wang in our university for technique help in gas chromatography. We also thank Professor Hongjun Wang at the Medical University of South Carolina, USA, for the helpful discussion during the writing process and Ms Wenjing Gou’s language assistance in preparing this manuscript.

## Author contributions

YR, ML, and KG conceived the experiments. YR, LL, ML, YW, and BW conducted the experiments. YR and KG analysed data. All authors were involved in writing the paper and had final approval of the submitted and published versions.

## Data availability

The manuscript and supplementary information files include all data and original Western blots.

## Supplementary data

This manuscript contains supplementary data.

## Competing interests

The authors declared no conflict of interest.

## Ethics approval and consent to participate

All animal experiments followed the Committee for Experimental Animals of Yangzhou University. All applicable institutional and/or national guidelines for the care and use of animals were followed.

## Institutional Review Board Statement

The animal study protocol was approved by the Ethics Committee of Yangzhou University (protocol code NSFC2020-SYXY-20 and dated 25 March 2020).

## ARRIVE guidelines Statement

The study was reported in accordance with ARRIVE guidelines.

## Consent for publication

Not applicable

## Notes

### Competing Interest Statement

The authors have declared no competing interest.

### Summary of Updates

(1) re-organising the logical order in the text to keep the consistency between the abstract and results; (2) changing all bar graphs into scatter dot plot graphs to understand the data point variance and clustering better using the Prism software; (3) changing the data of Table 1 (Hormone changes) into the scatter dot plot graph to show the individual values well; (4) correcting some errors and misunderstanding words.

