## supplementary information file for "Transgenic female mice producing *trans* 10, *cis* 12-conjugated linoleic acid present excessive prostaglandin E2, adrenaline, corticosterone, glucagon, and FGF21"

### Supplementary data

#### Supplementary Tables

Supplementary Table S1. Sources of dietary ingredients

| **Ingredients** | **Grams** | **Sources** |
|---|---:|---|
| Lactic casein | 200.0 | Gansu Hualing Dairy Co. Ltd, Hezuo city, China |
| L-cystine | 3.0 | Zhongtuan shihua Food Reagents Co. Ltd, Hangzhou, China |
| Sucrose | 176.8 | Jinhuitaiya, Chemcal Reagents Co. Ltd, Tianjin, China |
| Corn Starch | 452.2 | Zhucheng Xingmao Corn Co. Ltd, Weifang city, China |
| Maltodextrin 10 | 75.0 | Dongzhu Food Co. Ltd, Guilin city, China |
| Cellulose | 50.0 | Yisheng Kangyuan Biotech Co. Ltd, Zibo city, China |
| Soybean oil | 25.0 | Shandong Luhua Co. Ltd, Leiyang, China |
| Lard | 20.0 | Yaodong Baofengyuan Co. Ltd, Qingdao, China |
| Mineral Mix (20x) | 50.0 | Mingxin Chemical Reagents Co. Ltd, Zhenzhou, China |
| Choline bitartrate | 2.0 | Zhejiang Linuo Biotech Co. Ltd, Yiwu, China |
| Vitamin Mix (1000x) | 1.0 | Mingxin Chemical Reagents Co. Ltd, Zhenzhou, China |
| Total Gram | 1055 | |

**Supplementary Table S2. Primer sequences of real-time PCR**

| mRNA | Full name | Forward (top) and reverse (bottom) primers (5′ to 3′) | Genbank accession no. |
|---|---|---|---|
| *36B4* | Ribosomal protein, large, P0 | CACTGGTCTAGGACCCGAGAAG; GGTGCCTCTGGAGATTTTCG | NM_007475.5 |
| *Acaa2* | Acetyl-CoA acyltransferase 2 | CTGCTACGAGGTGTGTTCATC; AGCTCTGCATGACATTGCCC | NM_177470.3 |
| *Acbp* | Acyl-CoA-binding protein | CATCCGTATCACCTCACC; GTATTTACATCGCCCACA | NM_007830.4 |
| *Acox1* | Acyl-CoA oxidase 1, transcript variant 1 | AGATTGGTAGAAATTGCTGCAAAA; ACGCCACTTCCTTGCTCTTC | NM_015729.3 |
| *Acox2* | Acyl-CoA oxidase 2, transcript variant 1 | AACCCAGGGGATCGAGTGT; CGCAGCTCAGTGTTTGGGAT | NM_053115.2 |
| *Adcy3* | Adenylate cyclase 3, transcript variant 1 | AGATGTTCGGTGCCACCTG; ACTTCACCAGGGCTTCGTAAG | NM_138305.3 |
| | Adiponectin | AGCCGCTTATATGTATCGCTCA; TGCCGTCATAATGATTCTGTTGG | NM_009605.5 |
| *Agpat1* | 1-acylglycerol-3-phosphate O-acyltransferase 1 | GCTGGCTGGCAGGAATCAT; GTCTGAGCCACCTCGGACAT | NM_018862 |
| *Agpat2* | 1-acylglycerol-3-phosphate O-acyltransferase 2 | TTTGAGGTCAGCGGACAGAA; AGGATGCTCTGGTGATTAGAGATGA | NM_026212 |
| *Agrp* | Agouti related neuropeptide | GGCCTCAAGAAGACAACTGC; GACTCGTGCAGCCTTACACA | NM_007427.3 |
| *Ampk* | AMP-activated protein kinase | TTGACGATGAGGCTGTGAAG; ATAAGCCACTGCAAGCTGGT | NM_178143.2 |
| *ApoB* | Apolipoprotein B | CGTGGGCTCCAGCATTCTA; TCACCAGTCATTTCTGCCTTTG | NM_009693.2 |
| *Atgl* | Adipose triglyceride lipase | TTCACCATCCGCTTGTTGGAG; AGATGGTCACCCAATTTCCTC | NM_025802.3 |
| *Avp* | Arginine vasopressin | GCTCAACACTACGCTCTC; CTTGGGCAGTTCTGGAAG | NM_009732.2 |
| *Cart* | Cocaine- and amphetamine-regulated transcript | GTCCCACGAGAAGGAGCTGCCAA; GCCCATCCGCTCTCTGAGGGG | NM_013732.7 |
| *Cd11c* | CD11 antigen-like family member C, integrin alpha X | CTGGATAGCCTTTCTTCTGCTG; GCACACTGTGTCCGAACTC | NM_021334.3 |
| *Cd36* | CD36 antigen, transcript variant 1 | GGACATTGAGATTCTTTTCCTCTG; GCAAAGGCATTGGCTGGAAGAAC | NM_001159558.1 |
| *Cd68* | CD68 antigen | CTTCCCACAGGCAGCACAG; AATGATGAGAGGCAGCAAGAGG | NM_001291058.1 |

| | | | |
|---|---|---|---|
| *Cebpβ* | CCAAT/enhancer binding protein (C/EBP), beta | CTGCGGGGTTGTTGATGT; ATGCTCGAAACGGAAAAGGT | NM_001287738.1 |
| *Cgi-58* | Comparative gene identification 58 | TGGATTCTTGGCTGCTGCTTAC; TTAAAGGGAGTCAATGCTGCTC | NM_026179.2 |
| *Chrebp* | Carbohydrate response element binding protein | CCTTCGCCAACTCAGCACTT; TGGCTTGCTCAGGCACAA | NM_021455 |
| *Cpt1a* | Carnitine palmitoyltransferase 1a | CACCAACGGGCTCATCTTCTA; CAAAATGACCTAGCCTTCTATCGAA | NM_013495 |
| *Crh* | Corticotropin releasing hormone | GGCATCCTGAGAGAAGTCCC; GTTAGGGGCGCTCTCTTCTC | NM_205769.3 |
| *Cypb* | Cyclophilin B | TGGAGAGCACCAAGACAGACA; TGCCGGAGTCGACAATGAT | NM_011149.2 |
| *Deptor* | DEP domain containing MTOR-interacting protein | ATAGACGGCACCATCTCAAAAC; GTCGGCTAATTTCTGCATGAGT | NM_145470.3 |
| *Dgat1* | Diacylglycerol acyltransferase 1 | GAGGCCTCTCTGCCCCTATG; GCCCCTGGACAACACAGACT | NM_010046 |
| *Dgat2* | Diacylglycerol acyltransferase 2 | CCGCAAAGGCTTTGTGAAG; GGAATAAGTGGGAACCAGATCA | NM_026384 |
| *Dicer* | Ribonuclease type III | CACACGCCTCCTACCACTACAACAC; CCGTGGGTCTTCATAAAGGT | NM_148948.2 |
| *Dnmt1* | DNA (cytosine-5)-methyltransferase 1 | GTCGGACAGTGACACCCTTT; TGGGTTTCCGTTTAGTGGGG | NM_001199431.1 |
| *F4/80* | Adhesion G protein-coupled receptor E1 | CTTTGGCTAAGGGCTTCCAGTC; GCAAGGAGGACAGAGTTTATCGTG | NM_010130.4 |
| *Fads1* | Fatty acid desaturase 1 | AGCACATGCCATACAACCATC; TTTCCGCTGAACCACAAAATAGA | NM_146094.2 |
| *Fads2* | Fatty acid desaturase 2 | GATGGCTGCAACATGACTATGG; GCTGAGGCACCCTTTAAGTGG | NM_019699.2 |
| *Fasn* | Fatty acid synthase | GCTGCGGAAACTTCAGGAAAT; AGAGACGTGTCACTCCTGGACTT | NM_007988.3 |
| *Fgf21* | Fibroblast growth factor 21 | CTGCTGGGGGTCTACCAAG; CTGCGCCTACCACTGTTCC | NM_020013.4 |
| *Foxa2* | Forkhead box protein a2 | ACTTTGGGAGAGCTTTGAGGAA; CCCATCTATTTAGGGACACAGACA | NM_010446.3 |
| *Foxc2* | Forkhead box protein C2 | AAAGCGCCCCTCTCTCAGA;CTCAAAC TGAGCTGCGGATAAGT | NM_013519.2 |
| *G6p* | Glucose-6-phosphatase | TGGGCAAAATGGCAAGGA; TCTGCCCCAGGAATCAAAAAT | NM_008061.4 |
| *G6pd* | Glucose-6-phosphate dehydrogenase | GAACGCAAAGCTGAAGTGAGACT; TCATTACGCTTGCACTGTTGGT | NM_019468.2 |
| *Gapdh* | Glyceraldehyde-3-phosphate de- | GAACATCATCCCTGCATCC; CCAGTGAGCTTCCCGTTCA | NM_001289726.1 |

| | | | |
|---|---|---|---|
| | hydrogenase, transcript variant 1 | | |
| *Gck* | Glucokinase | CCGTGATCCGGGAAGAGAA; GGGAAACCTGACAGGGATGAG | NM_010292.5 |
| *Ghsr* | ghrelin receptor | GGACCAGAACCACAAACAGACA; CAGCAGAGGATGAAAGCAAACA | NM_177330.4 |
| *Glut4* | Glucose transporter type 4, transcript variant 1 | GTGACTGGAACACTGGTCCTA; CCAGCCACGTTGCATTGTAG | NM_009204.2 |
| *Gnrh* | Gonadotropin releasing hormone 1, transcript variant 1 | CACTGGTCCTATGGGTTGC; TTCTGCCTGGCTTCCTCT | NM_008145.3 |
| *Gpat1* | Glycerol-3-phosphate acyltrans-ferase 1, transcript variant 1 | CAACACCATCCCCGACATC; GTGACCTTCGATTATGCGATCA | NM_001356285.1 |
| *Grp78* | Glucose-regulated protein 78, transcript variant 1 | ACTTGGGGACCACCTATTCCT; ATCGCCAATCAGACGCTCC | NM_001163434.1 |
| *H19* | H19, transcript variant 1 | GGAATGTTGAAGGACTGAGGG; GTAACCGGGATGAATGTCTGG | NR_130973.1 |
| *Hmgcr* | HMG-CoA reductase, transcript variant 1 | CTTGTGGAATGCCTTGTGATTG; AGCCGAAGCAGCACATGAT | NM_008255.2 |
| *Hsl* | Hormone-sensitive lipase, transcript variant 1 | GGAGCACTACAAACGCAACGA; TCGGCCACCGGTAAAGAG | NM_010719.5 |
| *Htgl* | Hepatic triglyceride lipase, transcript variant 1 | ATGGGAAATCCCCTCCAAATCT; GTGCTGAGGTCTGAGACGA | NM_008280.2 |
| *Igf1* | Insulin-like growth factor 1, transcript variant 1 | GCCCCACTGAAGCCTACAAA; TGAGTCTTGGGCATGTCAGTGT | NM_010512.5 |
| *Igf2r* | Insulin-like growth factor 2 receptor | GCCTCTGGAGACATGAGGAC; GGTGCCTACTTTCCTGCTGA | NM_010515.2 |
| *Igfbp1* | Insulin-like growth factor binding protein 1 | ATCAGCCCATCCTGTGGAAC; TGCAGCTAATCTCTCTAGCACTT | NM_008341.4 |
| *Insig1* | Insulin induced gene 1 | TCACAGTGACTGAGCTTCAGCA; TCATCTTCATCACACCCAGGAC | NM_153526.5 |
| *Insig2a* | Insulin induced gene 2a, transcript variant 2 | CCCTCAATGAATGTACTGAAGGATT; TGTGAAGTGAAGCAGACCAATGT | NM_178082.3 |
| *Insr* | Insulin receptor, transcript variant 1 | CGAGTGCCCGTCTGGCTATA; GGCAGGGTCCCAGACATG | NM_010568.3 |
| *Irs1* | Insulin receptor substrate-1 | TCACAGCAGAATGAAGACC; CTACTGATGAGGAAGATATGAGG | NM_010570.4 |
| *Irs2* | Insulin receptor substrate-2 | GGAGAACCCAGACCCTAAGCTACT; GATGCCTTTGAGGCCTTCAC | NM_001081212.2 |
| *Lcad* | Long-chain acyl-CoA dehydrogenase | TCAATGGAAGCAAGGTGTTCA; GCCACGACGATCACGAGAT | NM_007381.4 |

| | | | |
|---|---|---|---|
| *Lchad* | Long-chain 3-hydroxyacyl-CoA dehydrogenase | TGCATTTGCCGCAGCTTTAC; GTTGGCCCAGATTTCGTTCA | NM_178878.3 |
| *Ldlr* | Low density lipoprotein receptor, transcript variant 1 | AGGCTGTGGGCTCCATAGG; TGCGGTCCAGGGTCATCT | NM_010700.3 |
| *Lepr* | Leptin receptor, transcript variant 1 | TGGTCCCAGCAGCTATGGT; ACCCAGAGAAGTTAGCACTGT | NM_146146.3 |
| *Lkb1* | Liver kinase B1, transcript variant 2 | GACTTCACAGTGCCTGGTGTC; AGCAGACAGGGAGCTACACTA | NM_001301853.2 |
| *Lpl* | Lipoprotein lipase | ACTATGTGTCTAACTGCCACTTCAA; ATACATTCCCGTTACCGTCCAT | NM_008509.2 |
| *Lxrα* | Liver X receptor α, transcript variant 1 | TACGTCTCCATCAACCACCCC; ACTTGCTCTGAATGGACGCTG | NM_013839.4 |
| *Lxrβ* | Liver X receptor β, transcript variant 1 | GCAGTTGGCACTAGAAG; GGTAGGCTGAGGTGTAA | NM_009473.3 |
| *Malic* | Malic enzyme 1 | GCCGGCTCTATCCTCCTTTG; TTTGTATGCATCTTGCACAATCTTT | NM_008615.2 |
| *Mc4r* | Melanocortic 4 receptor | CCCGGACGGAGGATGCTAT; TCGCCACGATCACTAGAATGT | NM_016977.4 |
| *Mcad* | Medium-chain acyl-CoA dehydrogenase | GCAACTGCCCGCAAGTTT; TACTCCCCGCTTTTGTCATATTC | NM_007382.5 |
| *Mgl* | Monoglyceride lipase, transcript variant 1 | CGGACTTCCAAGTTTTTGTCAGA; GCAGCCACTAGGATGGAGATG | NM_001166251.1 |
| *Npy* | Neuropeptide Y | CCTTCCATGTGGTGATGGGA; GCAGACTGGTTTCAGGGGAT | NM_023456.3 |
| *Npy1r* | Neuropeptide Y postsynaptic receptor Y1, transcript variant 1 | CACAGGCTGTCTTACACG; GCGAATGTATATCTTGAAGTAG | NM_010934.4 |
| *Nr3c1* | Nuclear receptor subfamily 3 group C member 1, transcript variant 1 | GGAAGCGTGATGGACTTGTAT; GCTTGGAATCTGCCTGAGAA | NM_008173.4 |
| *Orexins* | Orexins-total | GTCGCCAGAAGACGTGTTC; GGTGGTAGTTACGGTCGGAC | NM_010410.2 |
| *Oxt* | Oxytocin | TGGCTTACTGGCTCTGACCT; GGCAGGTAGTTCTCCTCCTG | NM_011025.4 |
| *Pai* | P*ropionibacterium acnes* isomerase | TGACGAGCGGGAATACTTTA; GAGGGTCATCAGCCCATCTA | |
| *Pepck* | Phosphoenolpyruvate carboxykinase | CCACAGCTGCTGCAGAACA; GAAGGGTCGCATGGCAAA | NM_011044.3 |
| *Pgc1α* | Ppar-γ coactivator 1 alpha, transcript variant 1 | CCCTGCCATTGTTAAGACC; TGCTGCTGTTCCTGTTTTC | NM_008904.3 |

| | | | |
|---|---|---|---|
| *Pgd* | 6-Phosphogluconate dehydrogenase, transcript variant 1 | TGAAGGGTCCTAAGGTGGTCC; CCGCCATAATTGAGGGTCCAG | NM_001081274.2 |
| *Pgr* | Progesterone receptor | CTCCGGGACCGAACAGAGT； ACAACAACCCTTTGGTAGCAG | NM_008829.2 |
| *Pi3k* | Phosphoinositide 3-kinase, transcript variant 2 | GACAGCGAAGCGACGGC; GTCTGATTTTACTGCCACGCTC | NM_001077495.2 |
| *Plin1* | Perilipin 1a | CTGTGTGCAATGCCTATGAGA; CTGGAGGGTATTGAAGAGCCG | NM_175640.2 |
| *Pomc* | Proopiomelanocortin, transcript variant 1 | CTCCTGCTTCAGACCTCCAT; CAGTCAGGGGCTGTTCATCT | NM_001278581.1 |
| *Ppar-γ* | Peroxisome proliferator-activated receptor-γ, transcript variant 1 | CACAATGCCATCAGGTTTGG; GCTGGTCGATATCACTGGAGATC | NM_001127330.2 |
| *Prdm16* | PR domain containing 16 | GACTTGGACACTACCACGGG; AGATGCACCCCCAAACTCAG | NM_027504.3 |
| *Scap* | SREBP cleavage-activating protein | ATTTGCTCACCGTGGAGATGTT; GAAGTCATCCAGGCCACTACTAATG | NM_001001144.3 |
| *Scd1* | Stearoyl-coA desaturase 1 | TCCTCCCTACCTCCAACT; CAACAACCAACCCTCGCA | NM_009127.4 |
| *Srebp1a* | Sterol regulatory element binding protein 1a, transcript variant X1/2 | GGCCGAGATGTGCGAACT; TTGTTGATGAGCTGGAGCATGT | NM_011480.4 |
| *Srebp1c* | Sterol regulatory element binding protein 1c, transcript variant X3 | GGAGCCATGGATTGCACATT; GGCCCGGGAAGTCACTGT | NM_001358314.1 |
| *Srebp2* | Sterol regulatory element binding protein 2 | GCGTTCTGGAGACCATGGA; ACAAAGTTGCTCTGAAAACAAATCA | NM_033218.1 |
| *Stat3* | Signal transducer and activator of transcription 3 | CTGTAGAGCCATACACCAAGCAGCAGC; GGTCTTCAGGTACGGGGCAGCAC | NM_213659.3 |
| *Ucp1* | Uncoupling protein 1 | GAGGTGTGGCAGTGTTCATTG; GGCTTGCATTCTGACCTTCA | NM_009463.3 |
| *Ucp2* | Uncoupling protein 2 | GCTTCTGCACCACCGTCAT; GCCCAAGGCAGAGTTCATGT | NM_011671.5 |

**Supplementary Table S3. Fatty acids (mg/g) in wild-type and Pai tissues**

| | Wild-type | Pai/wt | Pai/Pai |
|---|---|---|---|
| ***Hearts*** | | | |
| No. of samples | 5 | 4 | 4 |
| 8:0 | 0.010 ± 0.01 | 0.011 ± 0.00 | 0.010 ± 0.01 |
| 10:0 | ND | ND | ND |
| 12:0 | ND | ND | ND |
| 14:0 | 0.014 ± 0.00$^{a}$ | 0.015 ± 0.00$^{ab}$ | 0.019 ± 0.00$^{b}$ |
| 14:1 | ND | ND | ND |
| 16:0 | 1.364 ± 0.09$^{a}$ | 1.441 ± 0.20$^{ab}$ | 1.676 ± 0.21$^{b}$ |
| 16:1n-7 | 0.041 ± 0.01$^{a}$ | 0.056 ± 0.01$^{ab}$ | 0.052 ± 0.00$^{b}$ |
| 18:0 | 1.936 ± 0.25 | 1.803 ± 0.30 | 2.160 ± 0.21 |
| trans-18:1 | 0.016 ± 0.01 | 0.022 ± 0.01 | 0.019 ± 0.01 |
| 18:1n-9 | 0.760 ± 0.12$^{a}$ | 0.810 ± 0.17$^{ab}$ | 1.010 ± 0.08$^{b}$ |
| 18:1n-7 | 0.287 ± 0.03$^{a}$ | 0.328 ± 0.06$^{ab}$ | 0.355 ± 0.04$^{b}$ |
| 18:2n-6 | 1.880 ± 0.23 | 1.951 ± 0.38 | 2.120 ± 0.27 |
| 20:0 | 0.008 ± 0.00 | 0.009 ± 0.00 | 0.008 ± 0.00 |
| 18:3n-6 | ND | ND | ND |
| 18:3n-3 | 0.007 ± 0.00 | 0.009 ± 0.00 | 0.009 ± 0.00 |
| t10c12-CLA | ND | ND | ND |
| 20:3n-6 | 0.112 ± 0.03 | 0.113 ± 0.03 | 0.116 ± 0.01 |
| 20:4n-6 | 1.253 ± 0.31 | 1.256 ± 0.22 | 1.244 ± 0.16 |
| 20:5n-3 | 0.008 ± 0.00 | 0.008 ± 0.00 | 0.009 ± 0.00 |
| 24:1 | 0.053 ± 0.02 | 0.058 ± 0.00 | 0.058 ± 0.01 |
| 22:5n-3 | 0.142 ± 0.02 | 0.151 ± 0.03 | 0.155 ± 0.02 |
| 22:6n-3 | 2.673 ± 0.37 | 2.559 ± 0.46 | 3.087 ± 0.34 |
| Total FAs | 11.037 ± 1.09 | 11.176 ± 1.76 | 12.750 ± 1.30 |
| SFAs | 3.310 ± 0.34 | 3.252 ± 0.50 | 3.843 ± 0.41 |
| Unsaturated FAs | 7.367 ± 0.69 | 7.441 ± 1.21 | 8.368 ± 0.90 |
| MuFAs | 1.174 ± 0.15$^{a}$ | 1.298 ± 0.25$^{ab}$ | 1.518 ± 0.12$^{b}$ |
| PUFAs | 6.104 ± 0.53 | 6.067 ± 0.97 | 6.770 ± 0.78 |
| n-6 | 2.831 ± 0.37 | 2.724 ± 0.48 | 3.259 ± 0.36 |
| n-3 | 3.245 ± 0.56 | 3.319 ± 0.62 | 3.480 ± 0.44 |
| n-6/n-3 | 1.175 ± 0.34 | 1.230 ± 0.21 | 1.068 ± 0.06 |

**Supplementary Table S3. Fatty acids (mg/g) in wild-type and Pai tissues**

| | **Wild-type** | **Pai/wt** | **Pai/Pai** |
|---|---|---|---|
| n7/(n7+16:0) | 0.193 ± 0.01 | 0.210 ± 0.02 | 0.196 ± 0.01 |
| n7/(n7+18:0) | 0.282 ± 0.01[a] | 0.309 ± 0.02[ab] | 0.319 ± 0.01[b] |
| (n-6-LA)/n-6 | 0.417 ± 0.03[ab] | 0.413 ± 0.01[a] | 0.391 ± 0.01[b] |
| *Livers* | | | |
| No. of samples | 7 | 5 | 5 |
| 8:0 | 0.013 ± 0.00[a] | 0.007 ± 0.00[b] | 0.012 ± 0.00[ab] |
| 10:0 | ND | ND | ND |
| 12:0 | ND | ND | ND |
| 14:0 | 0.019 ± 0.01 | 0.020 ± 0.00 | 0.016 ± 0.01 |
| 14:1 | 0.009 ± 0.00[ab] | 0.007 ± 0.00[a] | 0.009 ± 0.00[b] |
| 16:0 | 1.916 ± 0.30 | 1.806 ± 0.19 | 1.654 ± 0.36 |
| 16:1n-7 | 0.121 ± 0.04 | 0.098 ± 0.02 | 0.090 ± 0.06 |
| 18:0 | 1.785 ± 0.23[a] | 1.736 ± 0.38[ab] | 1.465 ± 0.16[b] |
| trans-18:1 | 0.023 ± 0.01 | 0.030 ± 0.02 | 0.020 ± 0.01 |
| 18:1n-9 | 1.440 ± 0.29 | 1.382 ± 0.10 | 1.261 ± 0.31 |
| 18:1n-7 | 0.301 ± 0.11 | 0.288 ± 0.02 | 0.221 ± 0.11 |
| 18:2n-6 | 1.350 ± 0.21[a] | 1.104 ± 0.21[ab] | 1.045 ± 0.19[b] |
| 20:0 | 0.008 ± 0.00 | 0.007 ± 0.00 | 0.008 ± 0.00 |
| 18:3n-6 | 0.019 ± 0.00 | 0.018 ± 0.00 | 0.017 ± 0.00 |
| 18:3n-3 | 0.011 ± 0.00 | 0.009 ± 0.00 | 0.009 ± 0.00 |
| t10c12-CLA | 0.003 ± 0.00 | 0.004 ± 0.00 | ND |
| 20:3n-6 | 0.238 ± 0.08[a] | 0.173 ± 0.04[ab] | 0.150 ± 0.04[b] |
| 20:4n-6 | 1.768 ± 0.27[a] | 1.624 ± 0.41[ab] | 1.463 ± 0.18[b] |
| 20:5n-3 | 0.030 ± 0.01[a] | 0.019 ± 0.01[b] | 0.022 ± 0.01[ab] |
| 24:1 | 0.027 ± 0.00[a] | 0.026 ± 0.01[ab] | 0.021 ± 0.00[b] |
| 22:5n-3 | 0.031 ± 0.01[a] | 0.025 ± 0.00[b] | 0.024 ± 0.01[ab] |
| 22:6n-3 | 0.949 ± 0.12 | 0.923 ± 0.17 | 0.752 ± 0.16 |
| Total FAs | 10.903 ± 0.93[a] | 10.668 ± 1.77[ab] | 9.116 ± 1.17[b] |
| SFAs | 3.740 ± 0.35[a] | 3.576 ± 0.53[ab] | 3.154 ± 0.43[b] |
| Unsaturated FAs | 6.460 ± 0.90[a] | 5.871 ± 0.69[ab] | 5.211 ± 0.81[b] |
| MuFAs | 1.945 ± 0.40 | 1.865 ± 0.12 | 1.646 ± 0.45 |
| PUFAs | 4.421 ± 0.57[a] | 3.934 ± 0.66[ab] | 3.502 ± 0.47[b] |
| n-6 | 3.375 ± 0.49[a] | 2.919 ± 0.53[ab] | 2.676 ± 0.32[b] |
| n-3 | 1.022 ± 0.13[a] | 0.974 ± 0.16[ab] | 0.807 ± 0.17[b] |

**Supplementary Table S3. Fatty acids (mg/g) in wild-type and Pai tissues**

| | Wild-type | Pai/wt | Pai/Pai |
|---|---|---|---|
| n-6/n-3 | 3.322 ± 0.45 | 3.010 ± 0.45 | 3.373 ± 0.43 |
| n7/(n7+16:0) | 0.178 ± 0.04 | 0.177 ± 0.02 | 0.152 ± 0.04 |
| n7/(n7+18:0) | 0.444 ± 0.05 | 0.448 ± 0.05 | 0.458 ± 0.06 |
| (n-6-LA)/n-6 | 0.600 ± 0.03 | 0.619 ± 0.05 | 0.611 ± 0.04 |
| *Kidneys* | | | |
| No. of samples | 5 | 6 | 4 |
| 8:0 | 0.013 ± 0.01[a] | 0.004 ± 0.00[b] | 0.016 ± 0.00[a] |
| 10:0 | ND | ND | ND |
| 12:0 | ND | ND | ND |
| 14:0 | 0.022 ± 0.00[a] | 0.045 ± 0.02[b] | 0.021 ± 0.00[a] |
| 14:1 | 0.007 ± 0.00 | 0.008 ± 0.00 | 0.008 ± 0.00 |
| 16:0 | 1.830 ± 0.25[a] | 2.654 ± 0.57[b] | 1.981 ± 0.13[a] |
| 16:1n-7 | 0.062 ± 0.02[a] | 0.175 ± 0.08[b] | 0.062 ± 0.01[a] |
| 18:0 | 1.703 ± 0.25 | 2.097 ± 0.35 | 1.814 ± 0.13 |
| trans-18:1 | 0.012 ± 0.00 | 0.008 ± 0.00 | 0.010 ± 0.00 |
| 18:1n-9 | 0.839 ± 0.18[a] | 1.287 ± 0.27[b] | 0.904 ± 0.05[a] |
| 18:1n-7 | 0.306 ± 0.06[a] | 0.488 ± 0.07[b] | 0.333 ± 0.02[a] |
| 18:2n-6 | 1.024 ± 0.10[a] | 1.394 ± 0.15[b] | 1.055 ± 0.08[a] |
| 20:0 | 0.013 ± 0.00 | 0.016 ± 0.00 | 0.014 ± 0.00 |
| 18:3n-6 | 0.009 ± 0.00[a] | 0.012 ± 0.00[b] | 0.010 ± 0.00[a] |
| 18:3n-3 | 0.008 ± 0.00[a] | 0.015 ± 0.00[b] | 0.007 ± 0.00[a] |
| t10c12-CLA | 0.006 ± 0.00[a] | 0.011 ± 0.00[b] | 0.006 ± 0.00[a] |
| 20:3n-6 | 0.108 ± 0.02[a] | 0.142 ± 0.02[b] | 0.099 ± 0.01[a] |
| 20:4n-6 | 2.226 ± 0.33[a] | 2.863 ± 0.50[b] | 2.319 ± 0.16[a] |
| 20:5n-3 | 0.022 ± 0.01[ab] | 0.031 ± 0.01[a] | 0.019 ± 0.00[b] |
| 24:1 | 0.051 ± 0.01 | 0.054 ± 0.01 | 0.051 ± 0.00 |
| 22:5n-3 | 0.269 ± 0.48[ab] | 0.066 ± 0.01[a] | 0.049 ± 0.01[b] |
| 22:6n-3 | 0.937 ± 0.50 | 1.405 ± 0.42 | 1.350 ± 0.13 |
| Total FAs | 10.15 ± 1.39[a] | 13.32 ± 1.69[b] | 11.19 ± 1.47[a] |
| SFAs | 3.576 ± 0.50[a] | 4.817 ± 0.61[b] | 3.845 ± 0.24[a] |
| Unsaturated FAs | 5.999 ± 0.82[a] | 8.096 ± 1.06[b] | 6.389 ± 0.40[a] |
| MuFAs | 1.283 ± 0.25[a] | 2.046 ± 0.37[b] | 1.384 ± 0.07[a] |
| PUFAs | 4.633 ± 0.57[a] | 5.973 ± 1.07[b] | 4.945 ± 0.35[ab] |
| n-6 | 3.367 ± 0.43[a] | 4.411 ± 0.65[b] | 3.483 ± 0.24[a] |

**Supplementary Table S3. Fatty acids (mg/g) in wild-type and Pai tissues**

| | Wild-type | Pai/wt | Pai/Pai |
|---|---|---|---|
| n-3 | 1.231 ± 0.17 | 1.516 ± 0.43 | 1.426 ± 0.13 |
| n-6/n-3 | 2.743 ± 0.21[a] | 2.995 ± 0.37[a] | 2.450 ± 0.14[b] |
| n7/(n7+16:0) | 0.167 ± 0.01[a] | 0.199 ± 0.02[b] | 0.166 ± 0.01[a] |
| n7/(n7+18:0) | 0.329 ± 0.04 | 0.381 ± 0.07 | 0.333 ± 0.01 |
| (n-6-LA)/n-6 | 0.695 ± 0.02 | 0.682 ± 0.02 | 0.697 ± 0.01 |
| ***Skeletal muscle*** | | | |
| No. of samples | 4 | 4 | 4 |
| 8:0 | 0.035 ± 0.01 | 0.031 ± 0.02 | 0.025 ± 0.02 |
| 10:0 | ND | ND | ND |
| 12:0 | 0.028 ± 0.01 | 0.014 ± 0.01 | 0.009 ± 0.00 |
| 14:0 | 0.039 ± 0.01 | 0.035 ± 0.02 | 0.169 ± 0.29 |
| 14:1 | ND | ND | ND |
| 16:0 | 1.200 ± 0.21 | 0.887 ± 0.48 | 0.942 ± 0.48 |
| 16:1n-7 | 0.117 ± 0.03[a] | 0.132 ± 0.10[ab] | 0.075 ± 0.02[b] |
| 18:0 | 0.723 ± 0.10 | 0.450 ± 0.24 | 0.582 ± 0.33 |
| trans-18:1 | ND | ND | ND |
| 18:1n-9 | 0.398 ± 0.07 | 0.412 ± 0.31 | 0.326 ± 0.21 |
| 18:1n-7 | 0.167 ± 0.03 | 0.119 ± 0.07 | 0.130 ± 0.07 |
| 18:2n-6 | 1.229 ± 0.25[a] | 1.048 ± 0.32[ab] | 0.687 ± 0.20[b] |
| 20:0 | ND | ND | ND |
| 18:3n-6 | ND | ND | ND |
| 18:3n-3 | ND | ND | ND |
| t10c12-CLA | ND | ND | ND |
| 20:3n-6 | ND | ND | ND |
| 20:4n-6 | 0.685 ± 0.08 | 0.482 ± 0.26 | 0.581 ± 0.45 |
| 20:5n-3 | ND | ND | ND |
| 24:1 | ND | ND | ND |
| 22:5n-3 | 0.078 ± 0.03 | 0.048 ± 0.02 | 0.054 ± 0.02 |
| 22:6n-3 | 0.944 ± 0.35 | 0.593 ± 0.25 | 0.705 ± 0.12 |
| Total FAs | 6.423 ± 1.17 | 5.043 ± 2.37 | 5.833 ± 2.25 |
| SFAs | 2.071 ± 0.34 | 1.458 ± 0.78 | 1.775 ± 0.75 |
| Unsaturated FAs | 3.624 ± 0.70 | 2.842 ± 1.26 | 2.589 ± 1.06 |
| MuFAs | 0.683 ± 0.13 | 0.663 ± 0.47 | 0.530 ± 0.29 |
| PUFAs | 2.941 ± 0.64 | 2.170 ± 0.80 | 2.048 ± 0.77 |

**Supplementary Table S3. Fatty acids (mg/g) in wild-type and Pai tissues**

| | Wild-type | Pai/wt | Pai/Pai |
|---|---|---|---|
| n-6 | 1.914 ± 0.26 | 1.530 ± 0.56 | 1.283 ± 0.64 |
| n-3 | 1.027 ± 0.38 | 0.641 ± 0.26 | 0.765 ± 0.14 |
| n-6/n-3 | 1.973 ± 0.41 | 2.472 ± 0.57 | 1.618 ± 0.49 |
| n7/(n7+16:0) | 0.192 ± 0.02[ab] | 0.213 ± 0.02[a] | 0.182 ± 0.02[b] |
| n7/(n7+18:0) | 0.354 ± 0.04 | 0.454 ± 0.08 | 0.351 ± 0.03 |
| (n-6-LA)/n-6 | 0.362 ± 0.05 | 0.305 ± 0.05 | 0.431 ± 0.09 |
| *BAT* | | | |
| No. of samples | 4 | 4 | 4 |
| 8:0 | 0.012 ± 0.00 | 0.011 ± 0.00 | ND |
| 10:0 | ND | ND | ND |
| 12:0 | 0.012 ± 0.00[a] | 0.011 ± 0.00[a] | 0.016 ± 0.00[b] |
| 14:0 | 0.168 ± 0.02 | 0.170 ± 0.02 | 0.216 ± 0.07 |
| 14:1 | 0.012 ± 0.00[ab] | 0.011 ± 0.00[a] | 0.016 ± 0.00[b] |
| 16:0 | 2.559 ± 0.21 | 2.713 ± 0.44 | 3.354 ± 1.36 |
| 16:1n-7 | 0.451 ± 0.09 | 0.452 ± 0.17 | 0.717 ± 0.19 |
| 18:0 | 1.372 ± 0.20 | 1.367 ± 0.39 | 1.371 ± 0.45 |
| trans-18:1 | 0.096 ± 0.05 | 0.093 ± 0.06 | 0.080 ± 0.05 |
| 18:1n-9 | 3.177 ± 0.90 | 2.763 ± 0.81 | 3.420 ± 1.33 |
| 18:1n-7 | 0.610 ± 0.17 | 0.612 ± 0.15 | 0.824 ± 0.29 |
| 18:2n-6 | 1.878 ± 0.33 | 1.606 ± 0.37 | 1.683 ± 0.40 |
| 20:0 | ND | ND | ND |
| 18:3n-6 | 0.027 ± 0.01 | 0.021 ± 0.00 | 0.024 ± 0.01 |
| 18:3n-3 | 0.033 ± 0.01 | 0.028 ± 0.01 | 0.035 ± 0.01 |
| t10c12-CLA | 0.022 ± 0.02 | 0.029 ± 0.01 | 0.030 ± 0.00 |
| 20:3n-6 | 0.062 ± 0.00 | 0.066 ± 0.01 | 0.064 ± 0.01 |
| 20:4n-6 | 1.121 ± 0.28 | 1.135 ± 0.30 | 1.058 ± 0.21 |
| 20:5n-3 | 0.015 ± 0.00 | 0.017 ± 0.00 | 0.020 ± 0.00 |
| 24:1 | 0.023 ± 0.00 | 0.025 ± 0.01 | 0.028 ± 0.01 |
| 22:5n-3 | 0.022 ± 0.00 | 0.023 ± 0.00 | 0.022 ± 0.01 |
| 22:6n-3 | 0.342 ± 0.03 | 0.354 ± 0.07 | 0.317 ± 0.09 |
| Total FAs | 15.205 ± 3.09 | 14.043 ± 3.81 | 15.710 ± 5.56 |
| SFAs | 4.122 ± 0.36 | 4.270 ± 0.81 | 4.957 ± 1.88 |
| Unsaturated FAs | 8.050 ± 1.78 | 7.353 ± 1.67 | 8.455 ± 2.57 |
| MuFAs | 4.421 ± 1.18 | 3.991 ± 1.17 | 5.127 ± 1.83 |

**Supplementary Table S3. Fatty acids (mg/g) in wild-type and Pai tissues**

| | Wild-type | Pai/wt | Pai/Pai |
|---|---|---|---|
| PUFAs | 3.550 ± 0.60 | 3.285 ± 0.76 | 3.258 ± 0.73 |
| n-6 | 3.089 ± 0.61 | 2.828 ± 0.68 | 2.829 ± 0.62 |
| n-3 | 0.412 ± 0.03 | 0.421 ± 0.08 | 0.395 ± 0.10 |
| n-6/n-3 | 7.586 ± 2.01 | 6.687 ± 0.58 | 7.256 ± 0.73 |
| n7/(n7+16:0) | 0.291 ± 0.05 | 0.279 ± 0.06 | 0.324 ± 0.06 |
| n7/(n7+18:0) | 0.693 ± 0.03 | 0.663 ± 0.10 | 0.711 ± 0.02 |
| (n-6-LA)/n-6 | 0.390 ± 0.02 | 0.432 ± 0.01 | 0.407 ± 0.02 |

Note: Tissues of wild-type, Pai/wt, and Pai/Pai mice at the age of 11 weeks are for analysis, and 19:0 is an internal standard. Each value represents the mean ± SD. ND, Not detected; LA, linoleic acid; FA, fatty acids. Total FAs are total amounts from 8:0 to 22:6n-3 compositions. SFAs (saturated) and MUFAs (monounsaturated) are calculated as 8:0 + 10:0 + 12:0 + 14:0 + 16:0 + 18:0 + 20:0 + 22:0 + 24:0, and 14:1 + 16:1n-7 + trans-18:1 + 18:1n-9 + 18:1n-7 + 20:1 + 22:1 + 24:1, respectively. PUFAs is calculated as n-6 + n-3 as well as n-6 and n-3 are calculated as 18:2n-6 + 18:3n-6 + 20:3n-6 + 20:4n-6, and 18:3n-3 + 20:5n-3 + 22:5n-3 + 22:6n-3, respectively. n-7/(n-7 + 16:0) and n-7/(n-7 + 18:0) are calculated as (16:1n-7 + 18:1n-7)/(16:1n-7 + 18:1n-7 + 16:0), and 18:1n-9/(18:1n-9 + 18:0), respectively. Different letters in superscript indicate $p < 0.05$.

Supplementary Table S4. Dissection analysis of wild-type and Pai mice

| | wt | Pai/wt | Pai/Pai |
|---|---|---|---|
| Number of mice | 21 | 17 | 13 |
| Bodyweight (g) | 21.6 ± 1.8 | 22.0 ± 1.6 | 22.1 ± 1.6 |
| Heart weight (%) | 0.49 ± 0.03$^{a}$ | 0.51 ± 0.03$^{ab}$ | 0.55 ± 0.08$^{b}$ |
| Liver weight (%) | 4.78 ± 0.61$^{a}$ | 4.96 ± 0.55$^{ab}$ | 5.30 ± 0.58$^{b}$ |
| Spleen weight (%) | 0.43 ± 0.08$^{a}$ | 0.43 ± 0.07$^{a}$ | 0.55 ± 0.09$^{b}$ |
| Kidney weight (%) | 1.22 ± 0.12 | 1.27 ± 0.10 | 1.18 ± 0.37 |
| Lung weight (%) | 0.65 ± 0.10 | 0.63 ± 0.09 | 0.70 ± 0.13 |
| Pancreas weight (%) | 0.56 ± 0.08 | 0.52 ± 0.07 | 0.59 ± 0.10 |
| Brain weight (%) | 1.44 ± 0.15 | 1.41 ± 0.16 | 1.37 ± 0.13 |
| Hypothalamus weight (%) | 0.04 ± 0.01 | 0.04 ± 0.02 | 0.05 ± 0.02 |
| Ovary weight (%) | 0.04 ± 0.01$^{a}$ | 0.05 ± 0.01$^{b}$ | 0.05 ± 0.01$^{b}$ |
| WAT weight (%) | 0.35 ± 0.14 | 0.34 ± 0.16 | 0.32 ± 0.14 |
| BAT weight (%) | 0.41 ± 0.10 | 0.43 ± 0.09 | 0.41 ± 0.09 |
| Small intestine length (cm) | 35.5 ± 1.9 | 35.6 ± 2.2 | 37.0 ± 2.7 |

Note: Mice at 11 weeks of age were killed in the non-fasted state, and their issue/organ weight as a percentage of slaughter body weight was calculated. BAT, interscapular brown adipose tissue; WAT, peri-gonadal white adipose tissue. Each value represents the mean ± SD. Different letters in superscript indicate $p < 0.05$.

**Supplementary Figures.**

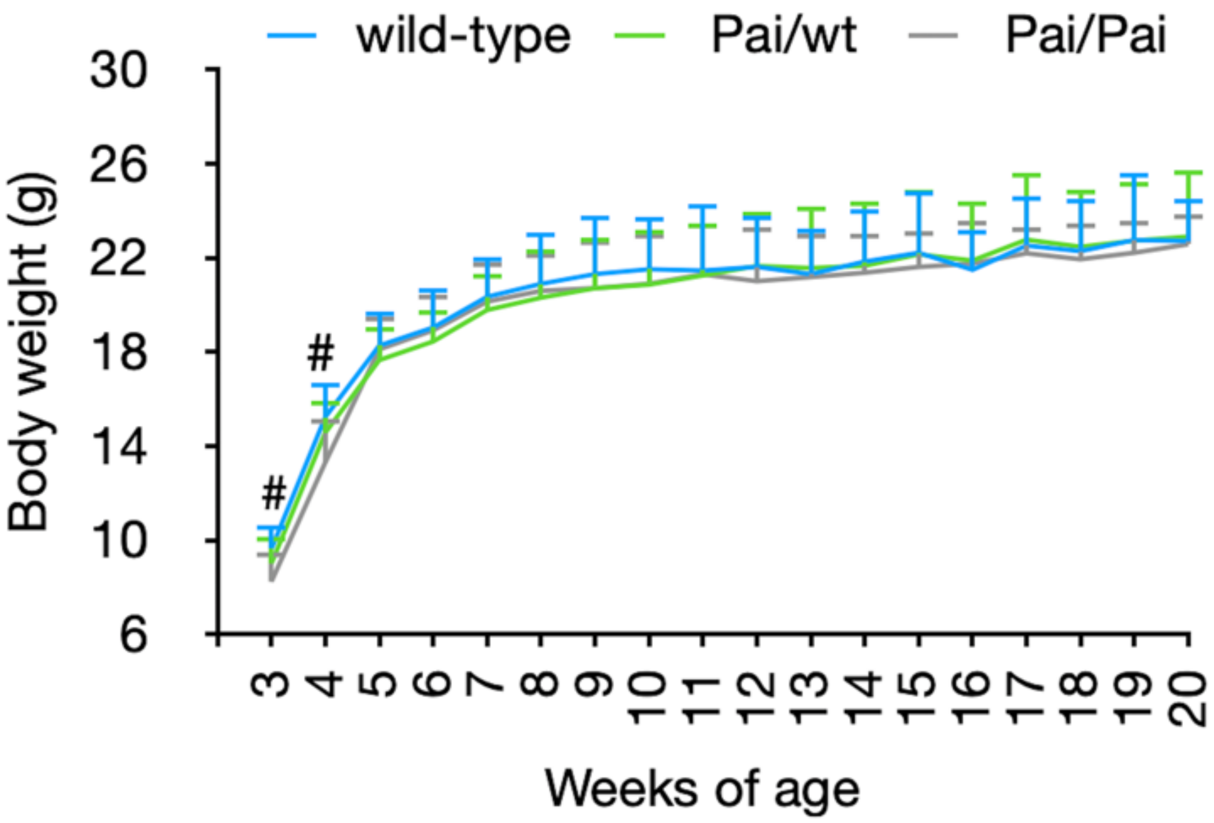


**Supplementary Figure S1. Comparison of growth of wild-type and Pai mice.** Each group contained 11-29 mice. Bars represent the mean ± SD. # indicates $p < 0.05$ among three genotypes.

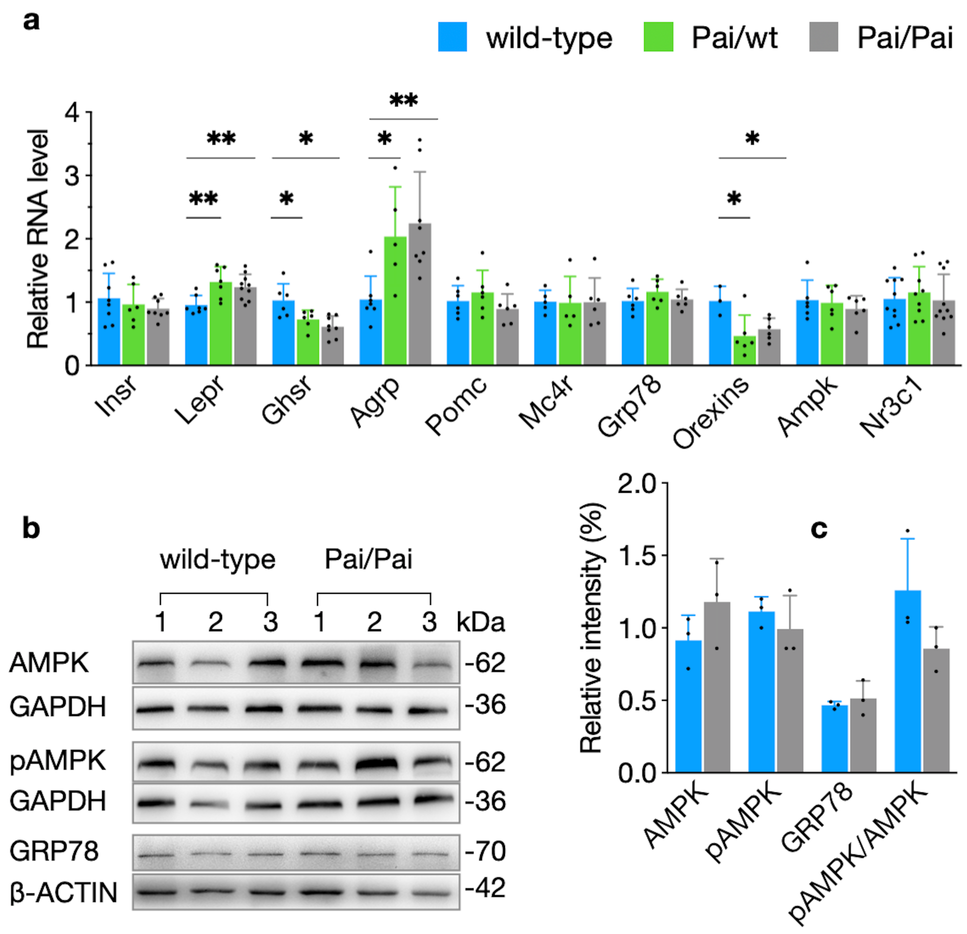


**Supplementary Figure S2. Analyses of the relative mRNAs (a), protein levels and their relative intensities in the hypothalamus (b-c).** Agrp, Agouti-related peptide; Ampk, AMP-activated protein kinase; Ghsr, Ghrelin receptor; Grp78, glucose-regulated protein 78; Insr, Insulin receptor; Lepr, Leptin receptor; Mc4r, Melanocortic 4 receptor; Nr3c1, Nuclear receptor subfamily 3 group C member 1; (p)AMPK, (phosphorylated) AMP-activated protein kinase; Pomc, Proopiomelanocortin. The bars represent the mean ± SD. * indicates $p < 0.05$ and ** indicates $p < 0.01$, respectively.

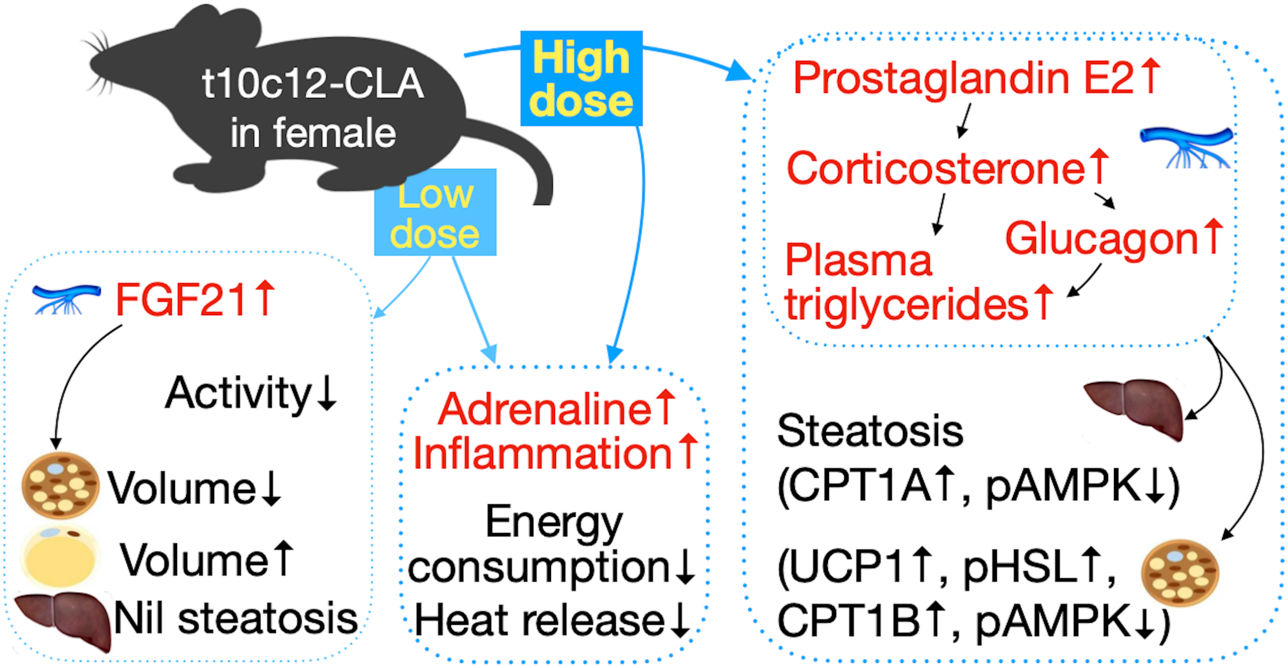


**Supplementary Figure S3. Diagram showing the dose-(in)dependent effect of chronic t10c12-CLA on lean female mice**. Besides the common (dose-independent) effects, such as increased adrenaline, inflammation reaction, less heat production, and low energy consumption (Middle panel), the low doses specifically increase circulating levels of FGF21, which plays an active role in maintaining energy homeostasis and suppress physical activity in Pai/wt mice (Left panel). Conversely, the high doses specifically result in excesses of prostaglandin E2, corticosterone, and glucagon; subsequently, the combination of these factors causes metabolic syndrome, such as fatty liver and hypertriglyceridemia through affecting the activities of critical enzymes involved in lipid metabolism in livers (CPT1A, pAMPK, etc.) and adipose tissues (UCP1, pHSL, CPT1B, pAMPK etc.) of Pai/Pai mice (Right panel). Words in black and red describe physiological features and endocrinal factors, respectively.
